# VCD-induced menopause mouse model reveals reprogramming of hepatic metabolism

**DOI:** 10.1101/2023.12.14.571644

**Authors:** Roshan Kumari, Michael E. Ponte, Edziu Franczak, John C. Prom, Maura F. O’Neil, Mihaela E. Sardiu, Andrew J. Lutkewitte, Kartik Shankar, E. Matthew Morris, John P. Thyfault

## Abstract

Menopause adversely impacts systemic energy metabolism and increases the risk of metabolic disease(s) including hepatic steatosis, but the mechanisms are largely unknown. Dosing female mice with vinyl cyclohexene dioxide (VCD) selectively causes follicular atresia in ovaries, leading to a murine menopause-like phenotype. In this study, we treated female C57BL6/J mice with VCD (160mg/kg i.p. for 20 consecutive days followed by verification of the lack of estrous cycling) to investigate changes in body composition, energy expenditure (EE), hepatic mitochondrial function, and hepatic steatosis across different dietary conditions. VCD treatment induced ovarian follicular loss and increased follicle-stimulating hormone (FSH) levels in female mice, mimicking a menopause-like phenotype. VCD treatment did not affect body composition, or EE in mice on a low-fat diet or in response to a short-term (1-week) high-fat, high sucrose diet (HFHS). However, the transition to a HFHS lowered cage activity in VCD mice. A chronic HFHS diet (16 weeks) significantly increased weight gain, fat mass, and hepatic steatosis in VCD-treated mice compared to HFHS-fed controls. In the liver, VCD mice showed suppressed hepatic mitochondrial respiration on LFD, while chronic HFHS diet resulted in compensatory increases in hepatic mitochondrial respiration. Also, liver RNA sequencing revealed that VCD promoted global upregulation of hepatic lipid/cholesterol synthesis pathways. Our findings suggest that the VCD-induced menopause model compromises hepatic mitochondrial function and lipid/cholesterol homeostasis that sets the stage for HFHS diet-induced steatosis while also increasing susceptibility to obesity.

## INTRODUCTION

Approximately 30% (∼ 50 million women) of the total US population are currently in or have gone through menopause, with this number projected to increase to 52 million by 2030^1^. Menopause is a natural biological process that occurs in women between the ages of 45-55 years, marking the end of menstrual cycles mainly due to hormonal changes that include a decline in circulating estrogen and an increase in follicle-stimulating hormone levels ^2, 3^. This hormonal shift, combined with natural physiological changes following menopause, has adverse effects on systemic energy metabolism and leads to a substantial increase in the prevalence of metabolic diseases such as obesity, dyslipidemia, insulin resistance, hepatic steatosis (excessive intrahepatic lipid storage), and type 2 diabetes (T2D) ^4–11^.

The prevalence of hepatic steatosis, which increases significantly following menopause, has attracted considerable attention in recent years. Hepatic steatosis, not due to excess alcohol consumption, has recently been re-coined “metabolic dysfunction associated-steatotic liver disease (MASLD)” by the American Association for the Study of Liver Disease but was previously labeled as non-alcoholic fatty liver disease. Hepatic steatosis, characterized by the accumulation of >5% lipid in the liver, contributes prominently to liver dysfunction and is often associated with obesity and insulin resistance ^12^. Understanding the relationship between menopause and risk for hepatic steatosis is crucial, given that women spend approximately 35% of their lives in the postmenopausal stage ^13–15^.

Decreased sex hormones after menopause increase the risk for steatosis and progression to steatohepatitis. Data from our group and others suggest the increased risk of MASLD may be linked to impaired mitochondrial function ^16, 17^. Further, increasing evidence indicates that hepatic mitochondrial dysfunction is a pathological hallmark in preclinical rodent models and human subjects with steatosis ^18–20^. It has been shown that during liver disease progression induced by obesity (humans) or high fat/high sucrose diets (HFHS) in rodents, there is an adaptive or compensatory response in hepatic mitochondria by increasing overall oxidative capacity. This is likely due to the excessive delivery of lipids and other nutrients to the liver, which increases the bioenergetic demand of hepatic mitochondria ^19, 21^. However, evidence in humans and rodents suggests that these compensatory adaptations to increase oxidative capacity drive oxidative stress, and collapse when liver injury progresses from steatosis to steatohepatitis and more pronounced liver injury ^22, 23^. Despite the clear clinical evidence that menopause drives risk for steatosis, there has been less focus on if and how menopause increases the risk for metabolic disorders and obesity via modulation of hepatic mitochondrial function. There is no doubt that the association between menopause and MASLD involves a complex interplay of hormonal changes, aging, and alterations in lipid metabolism ^24^. Although hormone replacement therapy has shown beneficial effects on metabolic outcomes in post-menopausal women ^25^, this and other therapeutic approaches (diet and exercise) would be strengthened by fully understanding the precise mechanisms by which menopause induces systemic and liver metabolic dysfunction ^26^.

Unlike women, mice do not go through natural menopause; instead, they go through estropause at varying times around 12 months of age ^27, 28^. Hence, the OVX model has been widely used as a blunt tool to induce a loss of ovarian function and somewhat mimic the effects of menopause on systemic and hepatic mitochondrial energy metabolism ^16^. However, OVX does not fully recapitulate human menopause, in which ovarian function declines over time and transitions from perimenopause to menopause and some residual ovarian hormone production remains. This limits the rigor and translatability of findings from the OVX model to menopause in women. A more physiologically relevant pre-clinical model using chronic administration of vinyl cyclohexene dioxide (VCD) has emerged to address this issue. VCD administration closely mimics human menopause by inducing a menopause-like phenotype (gradual follicular atresia, decreased estrogen, and increased follicle-stimulating hormone (FSH) in mice that maintain their ovaries and the synthesis of other ovarian hormones involved in regulating metabolism ^29–31^. The VCD model thus represents a valuable tool for understanding the impact of hormonal changes on energy metabolism, hepatic mitochondrial function, and steatosis. Therefore, the main goal of this study was to explore and characterize how VCD-induced menopause-like phenotype influences body composition, energy expenditure (EE), hepatic mitochondrial respiratory function, mitochondrial proteome, global hepatic gene expression, and the development of steatosis. Based on what occurs with menopause in women, we hypothesized that the VCD-induced menopausal model would reduce EE and metabolic flexibility, increase adiposity/obesity, alter hepatic mitochondrial function, and promote steatosis in mice. Overall, our findings suggest that VCD has pronounced effects on adiposity, hepatic gene expression for lipid and cholesterol synthesis, steatosis, and hepatic mitochondrial function but that these effects were most evident after chronic high-fat high sucrose diet intake.

## MATERIALS AND METHODS

### Animals, experimental design, and VCD-administration

All animal procedures were carried out in accordance with the guidelines for the Care and Use of Laboratory Animals published by the US National Institutes of Health on the humane treatment of experimental animals and with the approval of the Institutional Animal Care and Use Committee (IACUC) and Use Committee at the University of Kansas Medical Center. 20-22-weeks-old C57BL/6J female mice (#000664, Jackson Laboratory, Bar Harbor, ME, USA) were purchased and group housed at our facility at thermoneutrality (28⁰C) on 12:12 reverse light cycle (dark 10:00 – 22:00), with *ad libitum* access to water and standard chow diet. Mice were injected intraperitoneally with 160mg/kg BW of Viny cyclohexene dioxide (VCD, Sigma, 94956-100ML mixed in sesame oil) or vehicle (sesame oil, Sigma S3547) for 20 consecutive days. After injections, mice were individually housed with *ad libitum* access to a LFD (10% kcal fat, 3.5% kcal sucrose, 3.85 kcal/gram energy density, Research Diets, D12110704) until the confirmation of diestrus cycle 100 days after the first VCD injection occurred and after 10 days of continuous diestrus confirmation. The VCD mouse studies were divided into three separate experiments (#1, #2, and #3) which are outlined in Figure 1.

**Figure 1.**
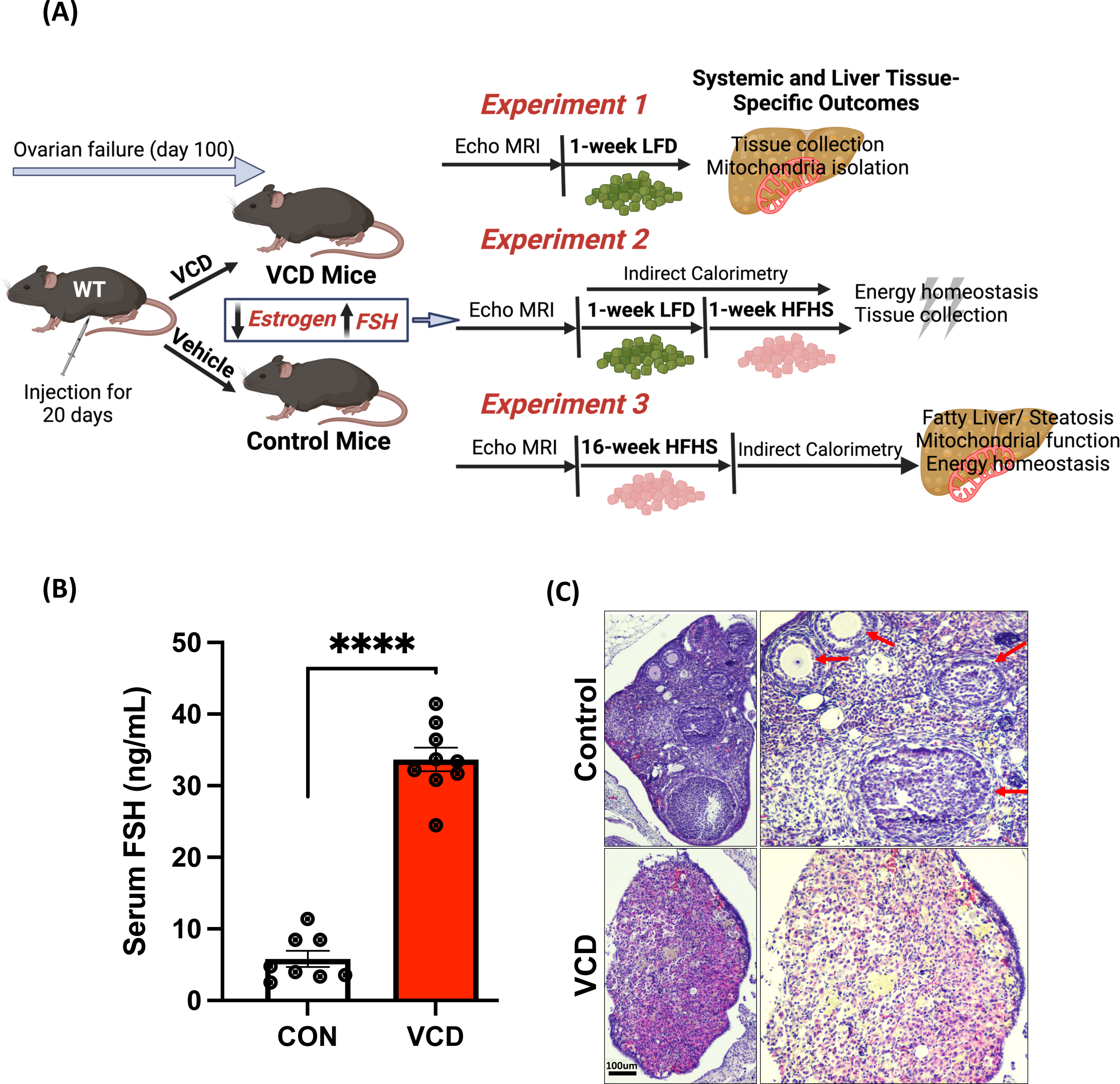
Working model and menopause confirmation. (A) Experimental design for each experiment (B) VCD administration showed significantly higher follicle stimulating hormone (FSH) levels in VCD treated mice compared to the control group, and (C) H&E staining representative image on the ovarian tissue sections; no sign of ovarian follicles in VCD treated mice (lower panel) compared to that of control mice (upper panel, showing clear ovarian follicles, red arrows, (scale bar: 100 μm). Data are presented as mean ± SEM (n=8-10/group), *p ≤ 0.05.

**Experiment #1:** After the confirmation of the diestrus cycle, VCD mice and control mice (n=8- 10/group) were fed a LFD for an additional week prior to sacrifice and analysis. Our first experiment aimed to validate the liver tissue-specific phenotype and mitochondrial outcomes in VCD treated mice compared to the control group (Figure 1A).

**Experiment #2:** Similar to experiment 1, after the confirmation of the diestrus cycle, VCD and control mice (n=8/treatment/diet) were fed additional one week of LFD and transitioned to HFHS for 1-week (45% kcal fat, 17% kcal sucrose, 4.73 kcal/gram energy density, Research Diets, D12451). The purpose of experiment 2 was to measure systemic energy metabolism via indirect calorimetry while on a LFD (1 week) and during the transition to a high-fat, high-sucrose diet (HFHS, D12451, 45% kcal fat, 17% kcal sucrose and 4.73 kcal/g energy density, Research Diets, New Brunswick, NJ, USA) for 1 week. Thus, experiment 2 followed the same experimental design as experiment 1 (Figure 1A) except for an additional 1-week on HFHS diet.

**Experiment #3:** After the confirmation of the diestrus cycle, VCD mice and control mice (n=8/group) were both fed HFHS for 16-weeks (45% kcal fat, 17% kcal sucrose, 4.73 kcal/gram energy density, Research Diets, D12451). This experiment focused on the effect of chronic HFHS on obesity development, steatosis, and mitochondrial measurements in VCD-treated mice and control mice.

### Anthropometrics and Energy Intake

Body weight measurement started on day 1 of VCD injection and was monitored weekly throughout the study. Food intake was assessed weekly. Body composition and food intake (fat mass and lean mass in grams) were measured after VCD induction and after both the one week and chronic HFHS feeding studies by EchoMRI-1100 system (EchoMRI, Houston, TX). Fat-free mass (FFM) was calculated as the difference between body weight (BW) and fat mass (FM). Energy intake was calculated as the energy density of LFD (3.85 kcal/g) or HFHS (4.73 kcal/gram) times the food intake for the 7 days.

### Indirect Calorimetry & Energy Metabolism

Promethion indirect calorimetry cages (Sable Systems International, Las Vegas, NV, USA), were utilized to assess energy metabolism in control and VCD mice from Experiment 2 for one week on LFD followed by one week on HFHS diet. Measures included total energy expenditure (TEE), resting energy expenditure (REE), respiratory quotient (RQ), and spontaneous cage activity as previously described ^32^. VCD mice and control mice (n=8/group) were housed individually in metabolic cages with ad *libitum* food and water access at 28°C. Mice were acclimated to the indirect cages for four days before data collection occurred. Energy expenditure (EE) per hour was calculated by a modified Weir equation [EE (kcal/hr) = 60*(0.003941*VO2+0.001106*VCO2)], and respiratory quotient (RQ) as VCO_2_/VO_2_. Weekly Total EE was calculated as the 7-day sum of the 24-hour EE for each day. Resting EE was determined as the EE (kcal/hr) during the 30-minute period with the lowest daily EE and summed across the 7 days. Activity EE is total EE minus resting EE and the thermic effect of food (data not shown). Thermic effect of food was determined from the consensus thermic effect of food for fat (2.5%), carbohydrate (7.5%), and protein (25%), and the manufacturer provided diet information for each diet ^32, 33^. As such, the thermic effect of food for HFHS (D12451, Research Diets, 4.73 kcal/g, 45% kcals fat, 35% kcal carbohydrate, 20% kcals protein) is 8.75% or 0.4139 kcal/g. Cage activity was calculated as all meters using the summed distances from Pythagoras’ theorem that the mouse moved based on XY second by second position coordinates.

### Tissue collection

Mice were euthanized using intraperitoneal injection of phenobarbital. Liver tissue was dissected, flash frozen in liquid nitrogen, and processed for RNA and protein while other portions were prepared for histology or mitochondrial isolation. At sacrifice, whole blood was collected via cardiac puncture followed by clotting at room temperature cooling on ice for 10 minutes and centrifuged at 4°C for 10 minutes at 8,000 x g for serum collection.

### Liver Triglycerides

25-30 mg of frozen powdered liver tissue was used to extract liver lipids as described previously ^34^ and reconstituted in a tert-butanol-Triton X solution (3:2) and liver triglycerides and cholesterol levels were measured via a commercially available kit (Sigma-Aldrich, St. Louis, MO).

### Histology

Liver tissue was fixed in 10% formalin at 4°C overnight, embedded in paraffin and sectioned at 5- 6 microns. After deparaffinizing, rehydrated sections were stained with hematoxylin and eosin (Cat. No. H-3502, Vector Laboratories) following the vendor’s protocol. Images were acquired using Nikon Auto Imaging System. Scoring was done by pathologist.

### NAS Scoring Criteria

The assessment of liver pathology involved grading according to the Nonalcoholic fatty liver disease activity score (NAS) criteria: Steatosis grades ranged from 0 (<5%), 1 (5-33%), 2 (33-66%), to 3 (>66%). The Lobular Inflammation grade included values of 0 for none, 1 for <2 foci/200x, 2 for 2-4 foci/200x, and 3 for >4 foci/200x. The Hepatocyte Ballooning grade encompasses 0 for none, 1 for few ballooned hepatocytes, and 2 for many/prominent ballooned hepatocytes.

### RNA Isolation and Gene Expression analysis

Total RNA was isolated from liver tissue using RNeasy Mini Kit (Qiagen, Germantown, MD) per manufacturer’s instructions from LFD and HFHS-fed VCD and control mice. cDNA was prepared according to the manufacturer’s protocol using the ImProm-II reverse transcription system (Promega, Madison, WI). Quantitation of gene expression was performed by real-time RT-PCR Quant Studio 3 (Thermo Fisher Scientific, Waltham, MA), using SYBR Green master mix (Applied Biosystems), normalized to *Cyclophilin B* in HFHS groups. The quantitative analysis was performed using the standard Λ1Λ1Ct method and results are shown as relative fold change in gene expression compared to control mice.

### RNA Seq and data analysis

RNA sequencing was performed in LFD-fed VCD mice and control mice (n=4). RNA was extracted from liver tissue as described previously. Isolated RNA from the liver was cleaned with the QIAamp RNA Blood Mini Kit (52304) according to the standard protocol. RNA-Seq was performed using the Illumina NovaSeq 6000 Sequencing System at the University of Kansas Medical Center – Genomics Core (Kansas City, KS). Quality control on RNA submissions was completed using the TapeStation 4200 using the RNA ScreenTape (Agilent Technologies 5067- 5576) prior to library preparation. Total RNA (1ug) was used to initiate the library preparation protocol. The total RNA fraction was processed by oligo dT bead capture of mRNA, fragmentation, reverse transcription into cDNA, end repair of cDNA, ligation with the appropriate Unique Dual Index (UDI) adaptors, strand selection and library amplification by PCR using the Universal Plus mRNA-seq with UDI preparation kit (Tecan Genomics 0520-A01).

Library validation was performed using the DNA 1000 ScreenTape (Agilent Technologies 5067- 5582) on the TapeStation 4200. The concentration of each library was determined by qPCR using the with the Roche Lightcycler96 using FastStart Essential DNA Green Master (Roche 06402712001) and KAPA Library Quant (Illumina) DNA Standards 1-6 (KAPA Biosystems KK4903). Libraries were pooled based on equal molar amounts to 1.9nM for multiplexed sequencing. Pooled libraries were denatured with 0.2N NaOH (0.04N final concentration) and neutralized with 400mM Tris-HCl pH 8.0. A dilution of the pooled libraries to 380 pM is performed in the sample tube, on instrument, followed by onboard clonal clustering of the patterned flow cell using the NovaSeq 6000 S1 Reagent Kit v1.5 (200 cycle) (Illumina 20028318). A 2x101 cycle sequencing profile with dual index reads is completed using the following sequence profile: Read 1 – 101 cycles x Index Read 1 – 8 cycles x Index Read 2 – 8 cycles x Read 2 – 101 cycles. Following collection, sequence data is converted from .bcl file format to fastq file format using bcl2fastq software and de-multiplexed into individual sequences for data distribution using a secure FTP site or Illumina BaseSpace for further downstream analysis.

Following sequencing and demultiplexing, all reads were trimmed for adapters, filtered based on quality score, and aligned to the mouse genome (mm10) using the STAR aligner. Resulting read alignments for each sample were imported in Seqmonk for gene level quantification as counts mapping to annotated genes. Gene counts were imported into R (v4.05) and analysis of differential expression between groups was done using the limma-voom pipeline ^35^. Transcripts with expression below 1 count/million were excluded from further analysis. Differentially expressed genes between two pair-wise comparisons (VCD vs control in LFD fed mice) were determined based on a p-value < 0.05 and a fold change of ±1.5 fold. A z-score cutoff of >2 was considered to identify pathways and regulators. Analysis of differentially expressed genes (DEG) for enrichment of gene ontology (GO) was done using Enrichr ^36^ and Ingenuity Pathway Analysis (Qiagen) ^37^. Heatmaps and other plots were generated using Morpheus software (Broad Institute).

### Hepatic mitochondrial isolation, respiration and H_2_O_2_ emission

Hepatic mitochondrial isolation was performed as described previously ^27, 38^. Briefly, liver tissue was quickly excised and submerged in 8 mL cold mitochondrial isolation buffer (220 mM mannitol, 70 mM sucrose, 10 mM Tris, 1 mM EDTA, pH adjusted to 7.4 with KOH) and homogenized on ice with a Teflon pestle. Homogenates were centrifuged (4°C, 10 min, 1500 x g), strained, and the supernatant was centrifuged again (4°C, 10 min, 8,000 x g). The supernatant was discarded and the resulting mitochondrial pellet with resuspended with 6 mL mitochondrial isolation buffer following centrifugation (4°C, 10 min, 6,000 x g). The process was repeated, using 4 mL mitochondrial isolation buffer with the addition of 0.1% fatty acid free BSA and lower centrifugation speed (4°C, 10 min, 6,000 x g). The final mitochondrial pellet was resuspended in MiR05 mitochondrial respiration buffer (0.5 mM EGTA, 3 mM MgCl2, 60 mM KMES, 20 mM glucose, 10 mM KH2PO4, 20 mM HEPES, 110 mM sucrose, 0.1% BSA, pH 7.1) and used in mitochondrial respiration studies. Mitochondrial respiration and H_2_O2 emission were concurrently assessed using Oroboros O2K fluorometer (Oroboros Instruments, Austria) as previously described ^27, 38^. All mitochondrial respiration experiments were performed with the starting substrates of 2mM malate, 10μM coenzyme A, and 2.5mM L-carnitine. To assess potential substrate differences in modulating mitochondrial respiration, two independent protocols were used with either potassium pyruvate (Pyr; 5mM) or palmitoyl-carnitine (PCoA; 10μM) added to the chambers to achieve basal respiration rates (Leak). Maximal coupled respiratory rate (State 3) was then determined following addition of 2.5mM adenosine 5’-diphosphate (ADP). To assess differences in CPT1a transport capacity, 10μM palmitoyl-carnitine (PC) was added to the chamber in PCoA supported respiration. For Pyr supported respiration, 2mM glutamate was added to assess further complex I driven respiratory rates. 10mM succinate was added to both protocols to determine maximal respiratory rates for ADP+succinate respiration (State 3S). Maximal uncoupled respiration was determined by tritrations of 0.1μM carbonyl cyanide-p-trifluoromethyoxyphenylhydrazone (FCCP; uncoupled) for each protocol. All data were normalized to total protein content of the isolated mitochondrial fractions added to the chamber, as determined by bicinchoninic acid protein assay (ThermoFisher Scientific, Waltham, MA). The coupling control ratio is calculated as the State 3 respiration divided by leak. ADP-dependent represents State 3 oxygen consumption minus leak. Data obtained from O2K were processed and analyzed using DatLab 7 Software. Data were normalized to mitochondrial protein content by the BCA method.

### Mitochondrial proteomics

100ug of mitochondria from each animal (n=8 each group) was spun down at 12000 g for 10 minutes at 4°C in 100uL of nuclease free water. Supernatant was discarded and dried mitochondrial pellets were collected for proteomics. Total protein from each sample was reduced, alkylated, and digested using single-pot, solid-phase-enhanced sample preparation with sequencing grade modified porcine trypsin (Promega) ^39^. Tryptic peptides were then separated by reverse phase XSelect CSH C18 2.5 um resin (Waters) on an in-line 150 x 0.075 mm column using an UltiMate 3000 RSLCnano system (Thermo). Peptides were eluted using a 60 min gradient from 98:2 to 65:35 buffer A:B ratio. Eluted peptides were ionized by electrospray (2.2 kV) followed by mass spectrometric analysis on an Orbitrap Exploris 480 mass spectrometer (Thermo). To assemble a chromatogram library, six gas-phase fractions were acquired on the Orbitrap Exploris with 4 m/z DIA spectra (4 m/z precursor isolation windows at 30,000 resolution, normalized AGC target 100%, maximum inject time 66 ms) using a staggered window pattern from narrow mass ranges using optimized window placements. Precursor spectra were acquired after each DIA duty cycle, spanning the m/z range of the gas-phase fraction (i.e., 496-602 m/z, 60,000 resolution, normalized AGC target 100%, maximum injection time 50 ms). For wide-window acquisitions, the Orbitrap Exploris was configured to acquire a precursor scan (385-1015 m/z, 60,000 resolution, normalized AGC target 100%, maximum injection time 50 ms) followed by 50x 12 m/z DIA spectra (12 m/z precursor isolation windows at 15,000 resolution, normalized AGC target 100%, maximum injection time 33 ms) using a staggered window pattern with optimized window placements. Precursor spectra were acquired after each DIA duty cycle. Buffer A = 0.1% formic acid, 0.5% acetonitrile, Buffer B = 0.1% formic acid, 99.9% acetonitrile.

### Mitochondrial proteomics data analysis

Following data acquisition, data were searched using an empirically corrected library against the UniProt *Mus musculus* database (January 2023) and a quantitative analysis was performed to obtain a comprehensive proteomic profile. Proteins were identified and quantified using EncyclopeDIA ^40^ and visualized with Scaffold DIA using 1% false discovery thresholds at both the protein and peptide levels. Protein exclusive intensity values were assessed for quality using ProteiNorm^41^. The data was normalized using cyclic loess^42^ and statistical analysis was performed using proteoDA^43^ with linear models for microarray data (limma) with empirical Bayes (eBayes) smoothing to the standard errors^42^. Proteins with an FDR adjusted p-value < 0.05 and a fold change > 2 were considered significant. All the proteins obtained from proteomics analysis were processed in Ingenuity Pathway Analysis (IPA) and identified up-and down-regulated proteins and the top pathways, proteins associated with the pathways and upstream regulators. These proteins were crossed-referenced with the Mitocarta dataset and examined for pathway enrichment among those proteins that were cross-referenced to Mitocarta^44^. Heatmaps were generated using Morpheus software, and the network was generated using the String database.

### Statistical Analysis

GraphPad Prism version 9.0 (GraphPad Software, San Diego, CA) was used for statistical analysis for each experiment unless otherwise mentioned. Mouse weights, body composition and energy data are expressed as mean ± standard error of the mean (SEM). Differences between CON vs. VCD were assessed via an independent *t-*test. A p-value < 0.05 was considered statistically significant for each experiment. Note: *p <0.05, **p < 0.005. The two-standard deviation test or ROUT method in Prism (GraphPad Software, San Diego, CA) was utilized to test for outliers. Two-way ANOVA was used to test for main effects and interactions of diet, and treatment for experiment 2. ANCOVA was performed in SPSS to determine the impact of differences in body weight on the effect of energy expenditure on 16-week HFHS diet between the two groups in experiment 3.

## RESULTS

### VCD administration promoted ovarian follicular loss and increased FSH levels in female mice

The overall study design and experimental groups are depicted in Figure 1A. Our initial phenotyping of the VCD treatment showed arrest of estrus cycling and ovarian failure as the persistent diestrus cycle of continuous 10 days with the majority of leukocytes by vaginal cytology (Supplementary Figure 1A). Additionally, we determined a menopause-like phenotype by measuring FSH levels of serum, VCD treated mice displayed significantly elevated FSH levels compared to control groups (Figure 1B). Also, H & E staining of ovarian sections of VCD-treated mice showed no sign of primordial or primary ovarian follicles (follicular atresia) in VCD-treated mice which were present in the control group shown by the red arrow (Figure 1C).

### VCD treatment on low-fat diet did not impact body mass/composition or energy intake

In experiment 1, we investigated the effects of VCD treatment on body weight, body composition, and weekly energy intake in mice fed a LFD. No substantial differences were observed in body weight, fat mass, fat free mass or energy intake between the VCD-treated and control groups. However, triglyceride levels in the liver of VCD-treated mice trended higher than those in the control group, though this difference didn’t reach statistical significance (Table 1). These results suggest that body weight and body composition remained relatively unchanged immediately after the onset of menopause in these female mice.

**Table 1.** Experiment 1: Low-fat diet (LFD) anthropometrics

|  |  | Control (Sesame Oil) | VCD |
| --- | --- | --- | --- |
| **Initial:** |  |  |  |
|  | Body weight (g) | 23.50 ± 0.55 | 23.76 ± 0.48 |
| **Final:** |  |  |  |
|  | Body weight (g) | 24.40 ± 0.38 | 25.34 ± 0.42 |
|  | Fat mass (g) | 3.88 ± 0.21 | 4.40 ± 0.27 |
|  | Fat-free mass (g) | 18.51 ± 0.33 | 18.88 ± 0.34 |
|  | Weekly food intake (g) | 15.46 ± 0.50 | 15.34 ± 0.48 |
|  | Weekly energy intake (kcal) | 59.53 ± 1.94 | 59.08 ± 1.86 |
|  | Liver triglycerides (mg/g) | 1.03 ± 0.09 | 1.28 ± 0.11 |

Data are presented as mean ± standard error of the mean (SEM), n=8-10/group).

### VCD treatment upregulates hepatic lipid and cholesterol synthesis pathways

Next, we wanted to determine if there are any transcriptome changes in the livers of VCD mice. To assess this, we performed bulk mRNA-seq gene expression analysis on liver samples. Transcriptome analysis identified 190 differentially expressed genes (DEG), with 151 genes downregulated and 39 genes upregulated. The upregulated genes were primarily involved in *de novo* lipogenesis, cholesterol and sterol biosynthetic metabolic pathways, as determined by Go- ontology analysis (Figure 2A). Furthermore, the downregulated genes were associated with cytokine regulation (Supplementary Figure 2A). A heatmap shows the upregulated genes (Figure 2B) and highlights genes involved in lipid and cholesterol pathways (Figure 2C). Significant changes were observed in key genes such as (*Fasn*, *Elovl6*, *Scd1*, *Acaca*, *Acacb*, *Acly*, and *Cyp8b1* in the liver of VCD mice compared to the control group, as identified by counts per million (CPM) values with unadjusted p value<0.05 from limma analysis (Supplementary Figure 2B). Although, H&E staining of liver sections showed no sign of steatosis in the liver sections of VCD mice relative to control mice on LFD (intact liver) (Supplementary Figure 1B). To dig deeper into the lipid-cholesterol metabolism pathway, upstream interaction analysis of these pathways and their associated DEG was performed using Ingenuity Pathway Analysis (IPA) software. This approach revealed the involvement of upstream regulators in the cholesterol pathway (Figure 2D). We also examined the expression of cholesterol transport proteins (StarD). Expression of *StarD9* was significantly downregulated and StarD4, an essential sterol transport protein involved in maintaining cholesterol homeostasis from cytoplasm to endoplasmic reticulum, trended to be higher (Supplementary Figure 3). Of note, expression of CfD (adipsin), a key gene involved in the alternative pathway of complement activation was decreased ∼30-fold.

**Figure 2.**
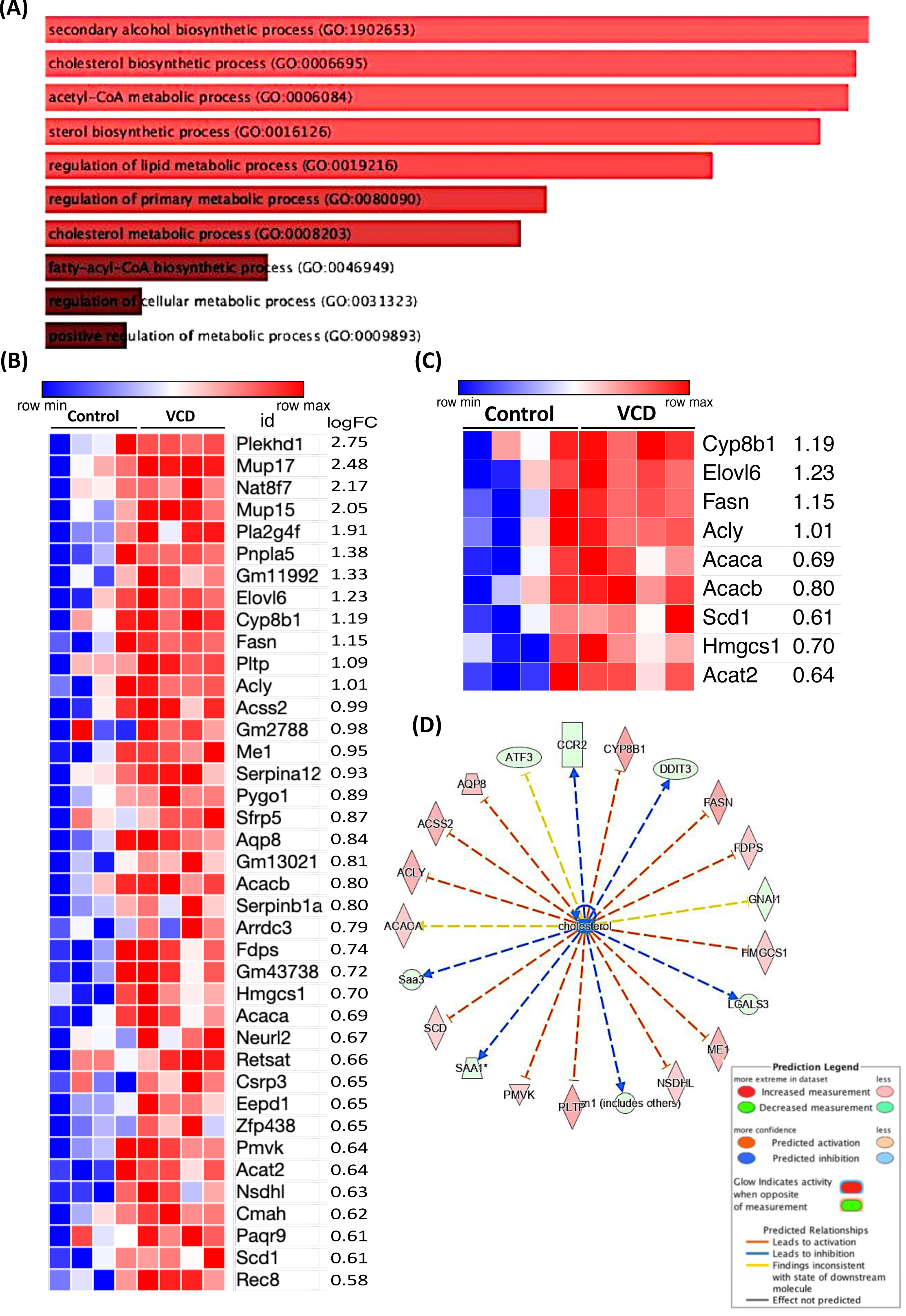
Experiment 1: VCD treatment upregulated lipid and cholesterol synthesis pathway. RNA Seq in VCD and control livers showing (A) Pathway (GO Ontology) upregulated in VCD, (B) Heatmap showing genes upregulated in VCD, (C) Heatmap showing lipid and cholesterol specific genes upregulated in VCD on LFD, and (D) Network analysis and associated DEGs (n=4/group), *p ≤ 0.05.

### VCD-treatment altered hepatic mitochondrial respiration and protein expression

Next, we aimed to determine if VCD treatment affected liver mitochondrial respiratory capacity. Consistent with previous findings in OVX mice ^16^, LFD fed VCD mice showed significantly lower leak, State 3, State 3S, and uncoupled hepatic mitochondrial respiration of fatty acid substrates in comparison to the control group (Figure 3A). We next leveraged proteomics in isolated hepatic mitochondria to explore possible mechanisms for loss of respiratory function induced by VCD. We focused on mitochondrial proteins associated with the electron transport chain, mitochondrial dynamics, including fission and fusion, and antioxidant proteins. The proteomics analysis identified 2562 total proteins. All proteins were cross-referenced to MitoCarta3.0 dataset as previously described ^44^ to identify only proteins within and associated with the mitochondria – a necessity as our isolated mitochondria contained both mitochondrial and non-mitochondrial proteins. This process identified 777 unique proteins found within the MitoCarta 3.0 dataset, 65 of the mitochondrial proteins had significant p-values between groups. In addition, 1771 non-mitochondrial proteins were differently expressed between groups, with 90 of the non-mitochondrial proteins exhibiting significant p-values (Figure 3B). The IPA analysis identified 20 significant pathways (Z-score = ± 2, -log *p*-value >1.3) with differential expression between the VCD-treated and control hepatic mitochondrial proteomes. Notably, pathways associated with mitochondrial respiration, particularly oxidative phosphorylation (z-score: -3.881), with 70 associated proteins, and the estrogen receptor signaling pathway (z-score: -2.309), involving 51 associated proteins were downregulated in VCD compared to controls (Figure 3C). A heatmap was generated to highlight all upregulated proteins specific to VCD cross-referenced with the MitoCarta dataset (Figure 3D). All MitoCarta proteins upregulated or downregulated, with statistically significant p-values and with logFC: ±1 is shown in (Supplementary Figure 4A-B). We also used Enrichr and REACTOME analysis on the MitoCarta derived proteins, revealing enriched pathways related to metabolism, the TCA cycle, mitochondrial transport, mitochondrial translation elongation, etc that were different between VCD and controls (Supplementary Figure 4C). Expanding our study to understand the connection between VCD treatment and hepatic mitochondrial quality, we performed IPA network analysis focused on mitophagy-associated regulators. The IPA analysis identified 18 proteins linked to mitochondrial autophagy (Z Score=- 1.666), including the downregulation of BNIP3 (a mitophagy protein), DNM1L, MFN1 and TOMM22 in VCD hepatic mitochondria compared to controls (Figure 3E). The protein-protein interaction network showed protein interactions for metabolic pathways generated by the string database (Supplementary Figure 4D. Overall, mitochondrial proteomics indicates a disruption in the regulation of mitochondrial homeostasis, including regulators of mitophagy, which were associated with suppressed mitochondrial respiration. All told, these data indicate VCD induced hepatic mitochondrial dysfunction prior to the onset of steatosis.

**Figure 3.**
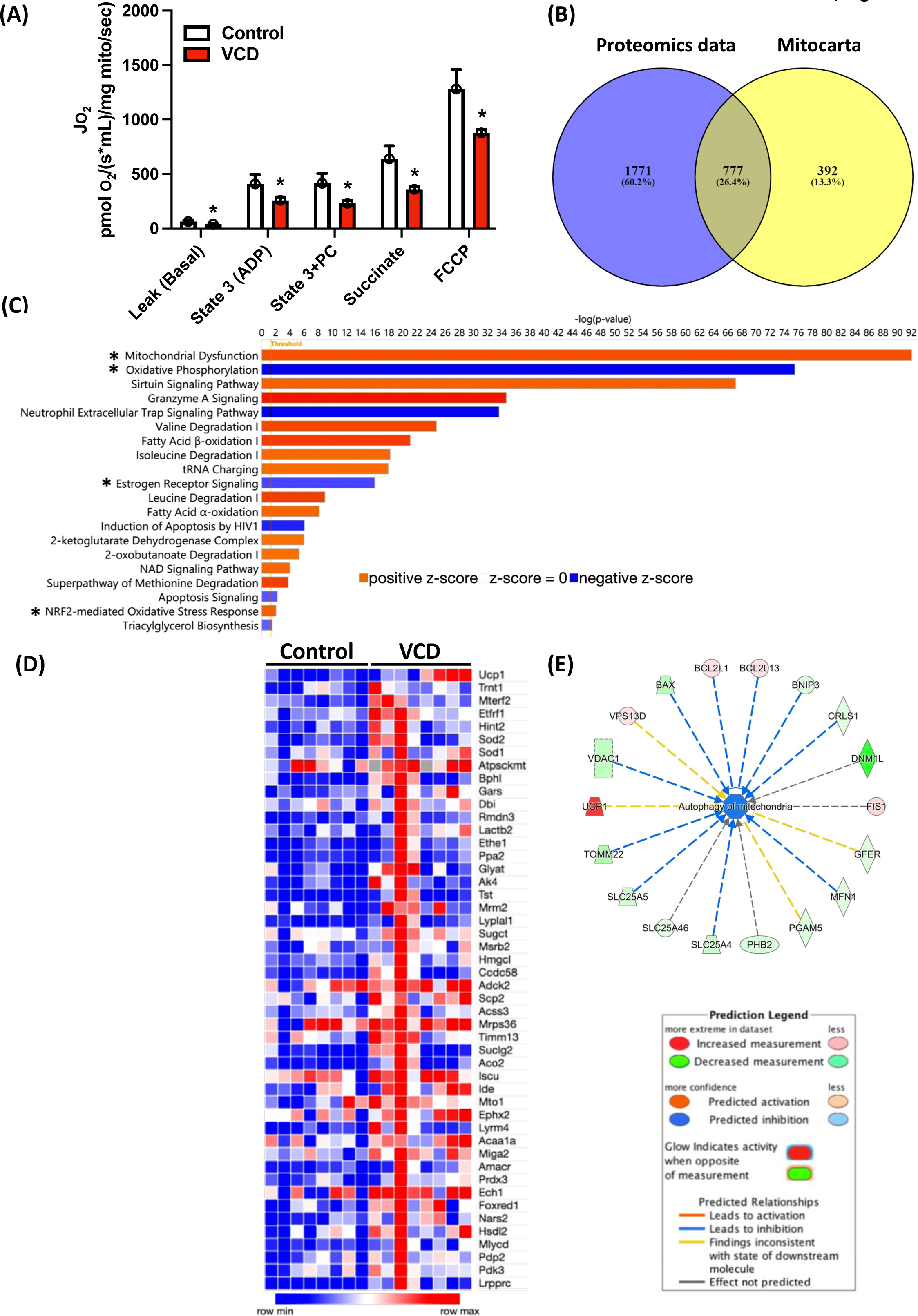
Experiment 1: VCD treatment alters mitochondrial respiratory capacity and proteomics pathways, primarily oxidative phosphorylation and the estrogen receptor signaling pathway. (A) Mitochondrial respiratory capacity was measured using palmitoyl-CoA (PC) as a main substrate at basal, ADP-dependent, Palmitoyl-CoA dependent, succinate- dependent, and FCCP uncoupled state from isolated liver mitochondria. Proteomics analysis in VCD hepatic mitochondria showing (B) Venn diagram showing Non-Mitocarta proteins in the VCD-treated group (1771), along with those proteins shared between (777) between Mitocarta and VCD, (C) Pathways affected by VCD treatment, (D) A heatmap showing upregulated proteins crossed referenced with Mitocarta with statistically significant p-values, and (E) Network analysis and regulators for autophagy of mitochondria. Respiration data was normalized to the mitochondrial protein via the BCA method. Data are presented as mean ± SEM (n=8-10), *p ≤ 0.05.

### VCD treatment did not impact liver injury markers or cholesterol levels on a LFD

To assess the effect of VCD treatment on liver injury markers and metabolic parameters, we measured alkaline phosphate (ALP) levels, aspartate aminotransferase (AST) and alanine amino transferase (ALT), triglycerides, glucose, and cholesterol levels in the serum from VCD and control on LFD diet in experiment 1 (Supplementary Table 1). No significant differences were observed for serum AST, ALT, triglycerides, or HDL and LDL cholesterol. However, VCD mice on LFD exhibited higher glucose levels (p=0.026) than controls.

### VCD treatment did not change energy expenditure in response to the 1-week HFHS diet

In experiment 2 we determined the effect of VCD treatment on basal energy metabolism by assessing EE during 1-week of LFD followed by 1-week of HFHS diet. We did not observe any differences in body weight, fat mass, and fat-free mass, total energy expenditure (TEE), resting EE, activity EE, or cage activity between the control and VCD-treated groups on LFD (Table 2). Transitioning to a 1-week HFHS diet also showed no differences in body weight, fat mass, fat-free mass, energy intake, TEE, or REE between the groups (Table 2). Activity EE and cage activity tended to be lower in VCD mice compared to control (p=0.08 & p=0.10, respectively). Next, we measured HFHS diet-induced changes in body weight, composition, energy expenditure, energy intake, and activity levels. The 1-Week HFHS feeding resulted in similar increases in body weight (Figure 4A), energy intake (Figure 4B), fat mass (Figure 4C), fat-free mass (Figure 4D), total EE (Figure 4E), and resting EE (Figure 4F) in both groups. Interestingly, VCD treated mass had a HFHS-induced reduction in cage activity (Figure 4G), with an associated trend towards reduced activity EE (Figure 4H) compared to control.

**Figure 4.**
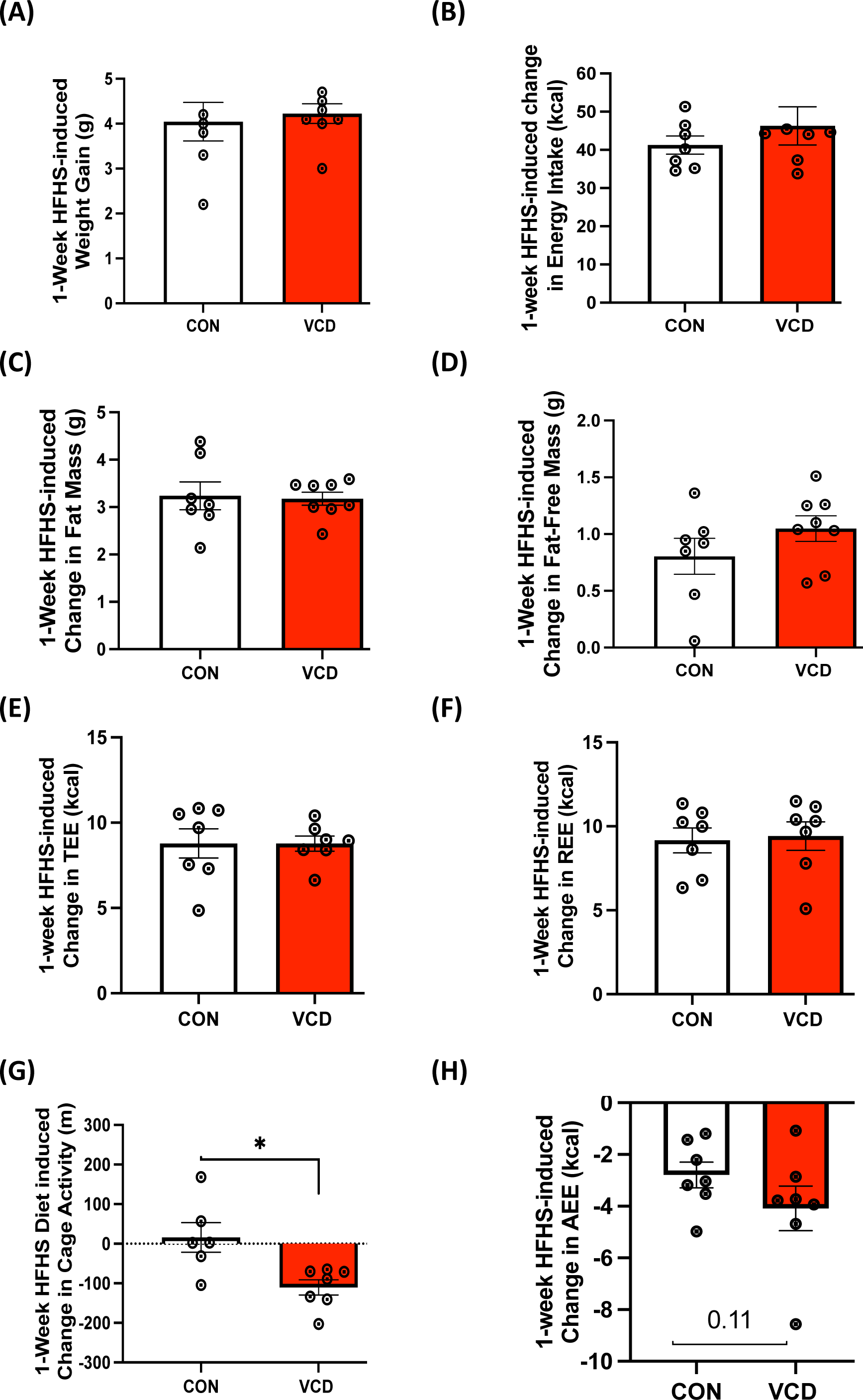
Experiment 2: Acute HFHS feeding reduced cage activity and activity EE in VCD mice. No differences in 1-week HFHS diet-induced change were observed in (A) weight gain, (B) energy intake), (C) fat mass, (D) fat-free mass, (E) total EE, or (F) resting EE. 1-week HFHS diet induced reductions in cage (G) cage activity and (H) activity EE in VCD mice compared to control. Values are shown as mean ± standard error of the mean (n=8/group).

**Table 2.**
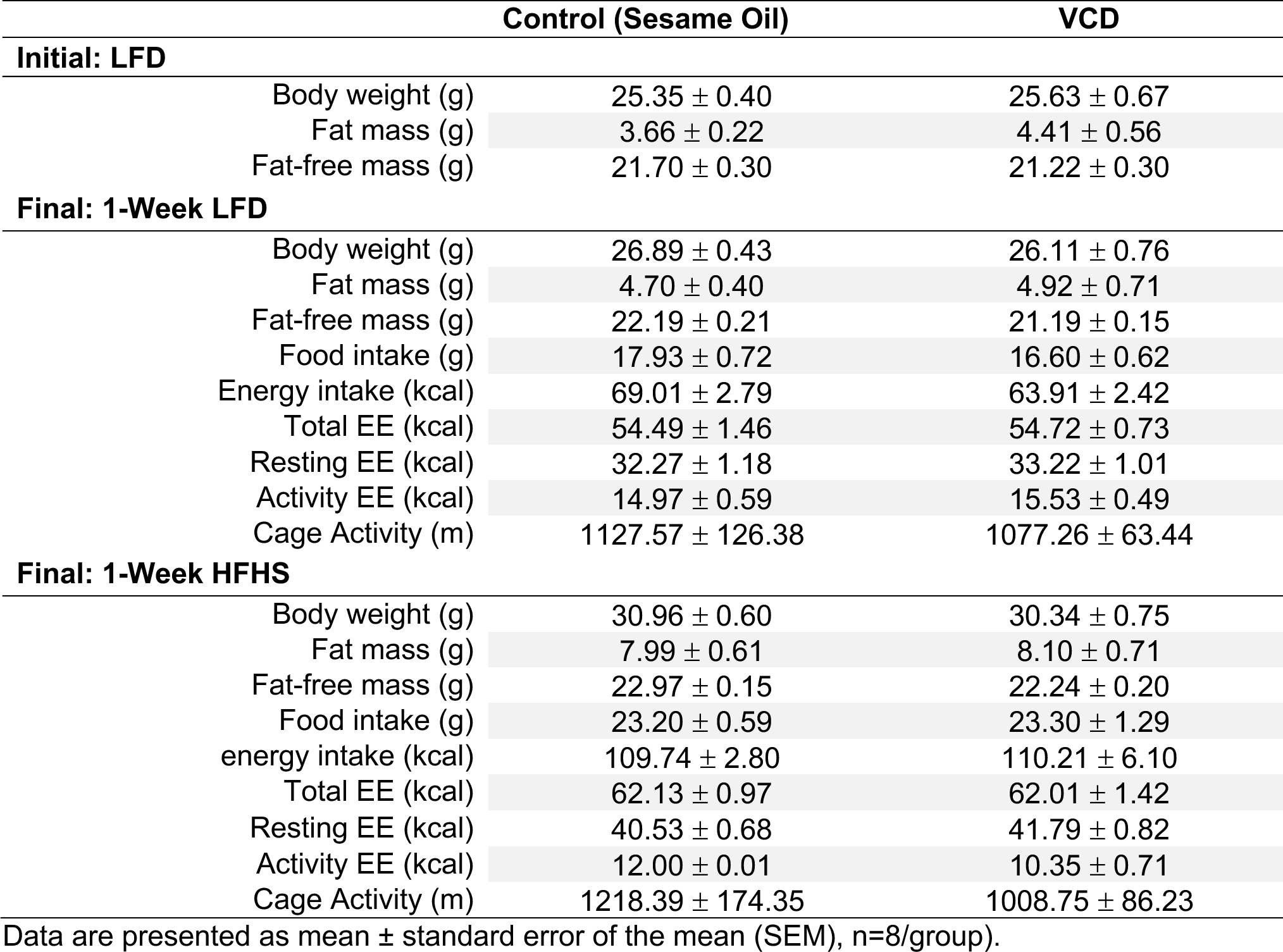
Experiment 2: Low-fat diet (LFD) and acute high-fat/ high-sucrose diet (HFHS) anthropometrics and energy metabolism

|  | Control (Sesame Oil) | VCD |
| --- | --- | --- |
| **Initial: LFD** |  |  |
| Body weight (g) | 25.35 ± 0.40 | 25.63 ± 0.67 |
| Fat mass (g) | 3.66 ± 0.22 | 4.41 ± 0.56 |
| Fat-free mass (g) | 21.70 ± 0.30 | 21.22 ± 0.30 |
| **Final: 1-Week LFD** |  |  |
| Body weight (g) | 26.89 ± 0.43 | 26.11 ± 0.76 |
| Fat mass (g) | 4.70 ± 0.40 | 4.92 ± 0.71 |
| Fat-free mass (g) | 22.19 ± 0.21 | 21.19 ± 0.15 |
| Food intake (g) | 17.93 ± 0.72 | 16.60 ± 0.62 |
| Energy intake (kcal) | 69.01 ± 2.79 | 63.91 ± 2.42 |
| Total EE (kcal) | 54.49 ± 1.46 | 54.72 ± 0.73 |
| Resting EE (kcal) | 32.27 ± 1.18 | 33.22 ± 1.01 |
| Activity EE (kcal) | 14.97 ± 0.59 | 15.53 ± 0.49 |
| Cage Activity (m) | 1127.57 ± 126.38 | 1077.26 ± 63.44 |
| **Final: 1-Week HFHS** |  |  |
| Body weight (g) | 30.96 ± 0.60 | 30.34 ± 0.75 |
| Fat mass (g) | 7.99 ± 0.61 | 8.10 ± 0.71 |
| Fat-free mass (g) | 22.97 ± 0.15 | 22.24 ± 0.20 |
| Food intake (g) | 23.20 ± 0.59 | 23.30 ± 1.29 |
| energy intake (kcal) | 109.74 ± 2.80 | 110.21 ± 6.10 |
| Total EE (kcal) | 62.13 ± 0.97 | 62.01 ± 1.42 |
| Resting EE (kcal) | 40.53 ± 0.68 | 41.79 ± 0.82 |
| Activity EE (kcal) | 12.00 ± 0.01 | 10.35 ± 0.71 |
| Cage Activity (m) | 1218.39 ± 174.35 | 1008.75 ± 86.23 |

Data are presented as mean ± standard error of the mean (SEM), n=8/group).

### VCD-treated mice exhibit reduced respiratory quotient

To determine whether there were any differences in substrate utilization between VCD and control mice at baseline or following the transition to a 1-week HFHS we quantified daily respiratory quotient (RQ). VCD mice had lower respiratory quotient during both LFD and HFHS feeding (Figure 5A & B). Further, VCD mice may be metabolically inflexible, as they tended to have a less of a reduction in RQ following the transition to HFHS compared to control (Figure 5C & D). These data demonstrate that VCD mice preferentially utilize lipids for energy metabolism regardless of dietary exposure and suggest that loss of follicular function impairs the capacity to alter systemic metabolism to increased dietary fat.

**Figure 5.**
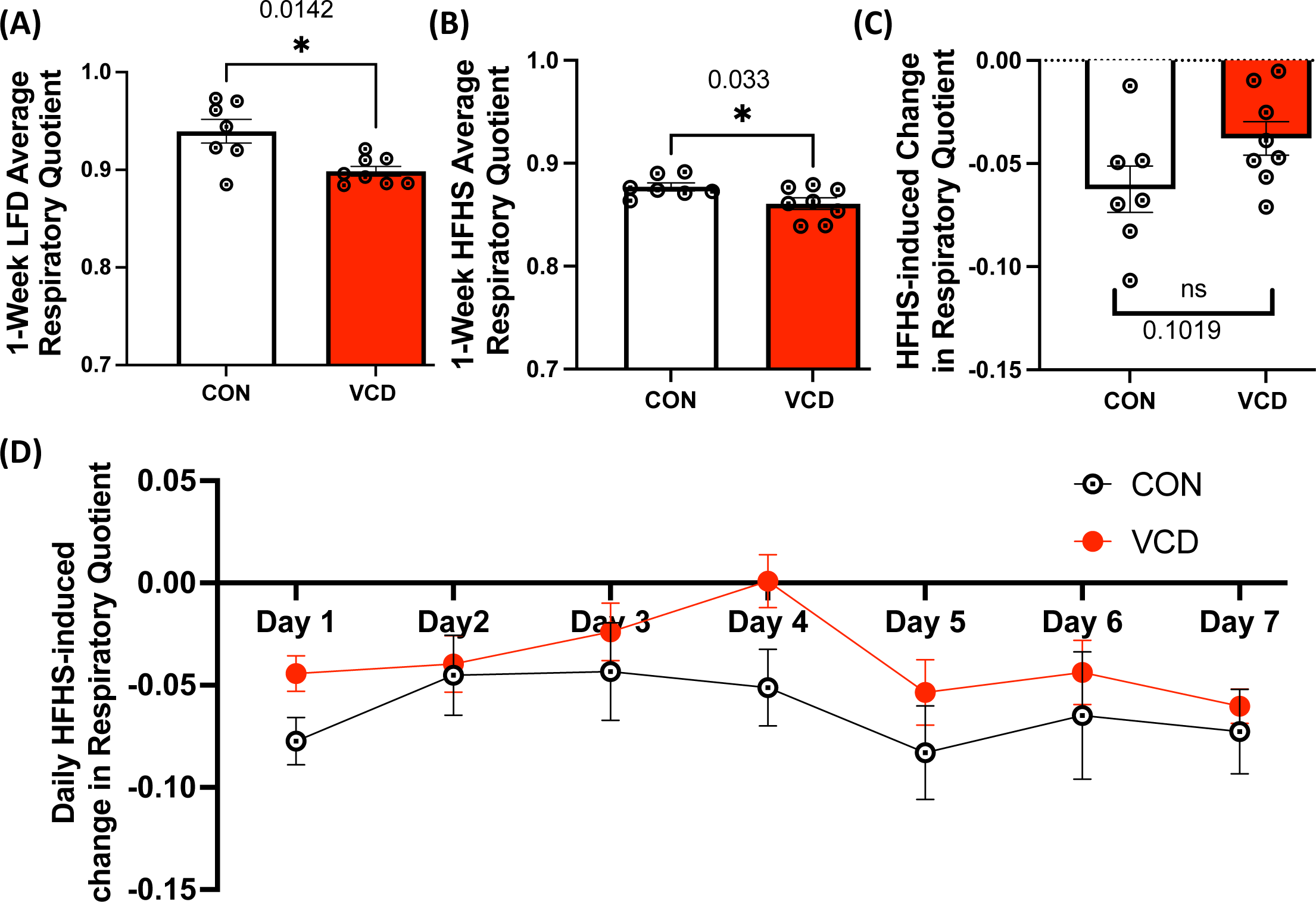
Experiment 2: VCD-treated mice exhibit increased fat utilization during LFD and acute HFHS feeding. VCD treated mice show (A) reduced respiratory quotient (RQ) on LFD, (B) reduced RQ during 1-week of HFHS feeding and tended to have reduced (C) HFHS-induced changes in RQ. (D) Daily RQ changes during HFHS exposure from LFD (baseline). Data are presented as mean ± SEM (n=8), *p ≤ 0.05.

### VCD mice showed greater susceptibility for short-term HFHS diet-induced steatosis and altered gene expression for hepatic lipogenesis and inflammation

To determine the effect of VCD treatment on liver lipid loads and steatosis in these females fed a 1-week HFHS diet, we examined liver tissue from both VCD treated and control mice. VCD treated mice displayed an increased trend in triglyceride levels on the HFHS diet, compared to control mice (Figure 6A). Also, an additional week of HFHS feeding led to steatosis in both groups, with VCD group showing evidence of larger lipid droplets (not quantified) (Figure 6B). We next wanted to perform a targeted examination for lipogenic, mitochondrial and inflammatory gene expression in the livers of VCD and control mice after the 1-week HFHS diet. As expected, the high-fat diet led to a reduction in lipogenesis. Livers from VCD mice had decreased expression of several genes involved in lipogenesis such as *Fasn*, *Acly*, *and Scd1* on the HFHS diet (Figure 6C-E). No changes in gene expression were observed with mitochondrial markers (Figure 6F-H). The expression of inflammatory markers *Tnfɑ* and *Erm1* did not show major differences, while IL6 trended up but was not significantly different from the control (Figure 6I-K). Taken together, this data suggests that acute HFHS diet led to a reduction in lipogenesis that was more pronounced in VCD than controls.

**Figure 6.**
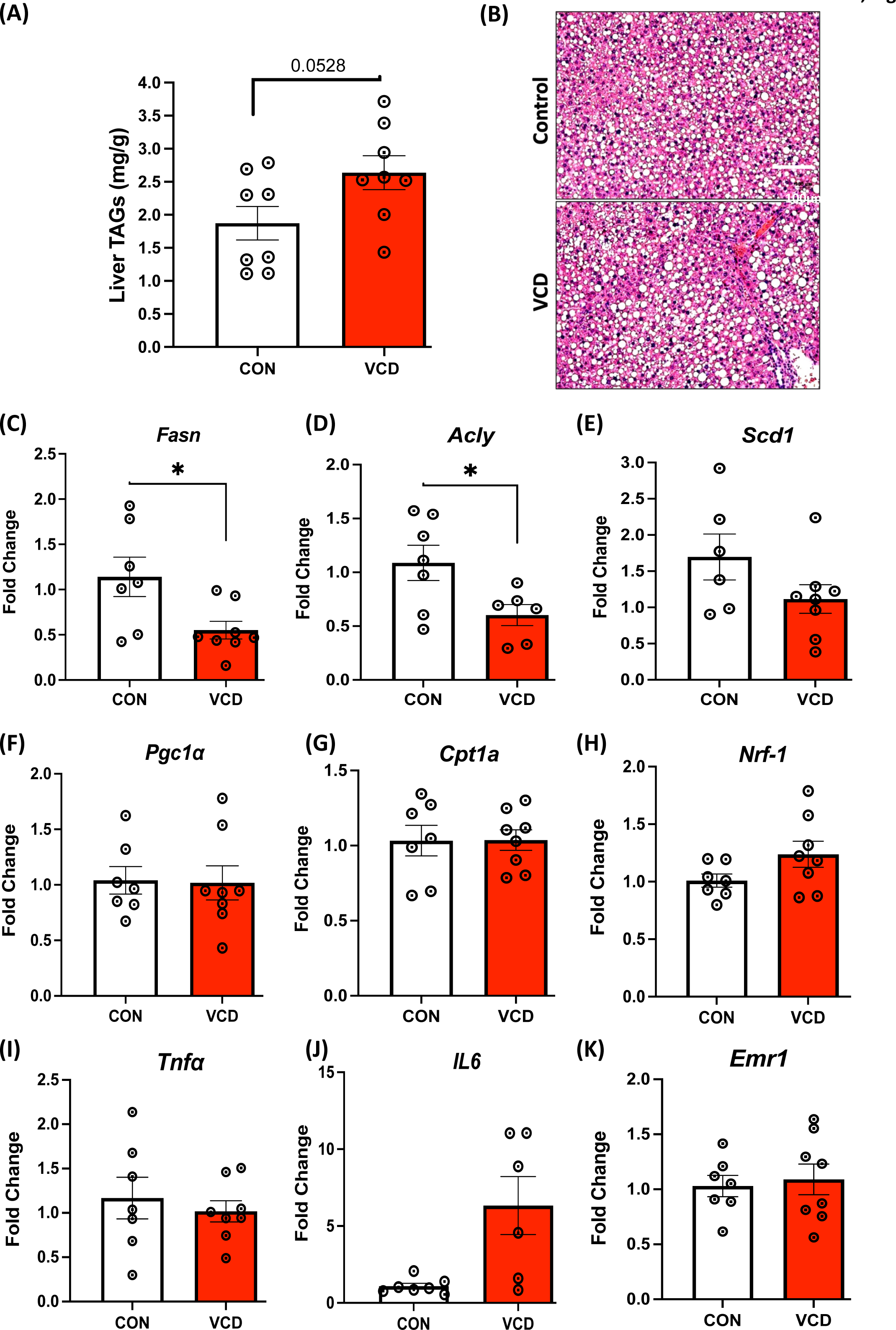
Experiment 2: Increased steatosis and reduced expression of key genes involved in lipogenesis in VCD livers during 1-week HFHS. (A) liver triglyceride content on acute HFHS diet, (B) Representative H & E staining of liver sections (scale bar: 100 μm). mRNA levels showing (C-E) Lower expression of *Fasn, Acly, and Scd1* in VCD treated liver samples compared to control mice on HFHS diet (n=7-8), (F-H) No change observed in mitochondrial markers, and (I-K) Inflammatory marker. Data are presented as mean ± SEM (n=7-8/group), *t*-tests were used to evaluate statistical significance, *p<0.05.

### VCD mice showed no evidence of liver injury or disrupted cholesterol homeostasis on short term HFHS diet

Similar to LFD, we assessed the effect of VCD treatment on liver injury markers on acute HFHS diet. Though VCD-treated mice tended to have higher ALP, AST or ALT levels compared to control groups, it didn’t reach statistically significant. No difference was found in triglycerides or glucose levels between the two groups (Supplementary Table 2). It is worth noting that higher VCD-induced fasting glucose levels on LFD (Supplementary Table 1), were no longer elevated following the 1-week HFHS diet. These data demonstrate that VCD treatment did not elicit any acute HFHS-induced alterations in liver injury markers, triglycerides, or cholesterol levels in the serum.

### VCD mice displayed greater adiposity and changes in energy expenditure on chronic HFHS diet

To assess the interaction of VCD treatment and chronic HFHS diet, we fed mice HFHS for 16- weeks following the confirmation of menopause (experiment 3) and monitored body weight, body composition, and other metabolic parameters. After 16-weeks, VCD treated mice showed significantly higher body weight with significant increase in fat mass compared to the control mice (Table 3). Also, weekly total EE, resting EE, and energy intake were greater in VCD following chronic HFHS feeding. To assess whether the observed differences in systemic energy metabolism between control mice and VCD mice are due to the difference in body weight and composition, we performed ANCOVA to generate adjusted marginal means with body weight and composition (fat mass + fat-free mass) as covariates. Following ANCOVA analysis, no difference for total EE, resting EE, or EI was observed between control and VCD mice. Interestingly, while body weight was the most powerful co-variate in this analysis, fat mass was a strong co-variate (as partial Eta-squared) for all the outcomes tested. Unlike following acute HFHS feeding, no differences were observed in RQ, cage activity, or activity EE between VCD and control mice during chronic HFHS feeding. These data demonstrate that differences in energy metabolism between VCD and control mice following chronic HFHS are due primarily to the increased weight gain and adiposity of VCD mice.

**Table 3.** Experiment 3: Chronic high-fat/high-sucrose diet (HFHS) anthropometrics and energy metabolism

| Anthropometrics |  |  |  |
| --- | --- | --- | --- |
| Initial: | Control (Sesame Oil) | VCD |  |
| Body weight (g) | 23.68 ± 0.30 | 23.06 ± 0.76 |  |
| Menopause (before HFHS) |  |  |  |
| Body weight (g) | 22.95 ± 0.37 | 24.29 ± 0.77 |  |
| Fat mass (g) | 2.54 ± 0.21 | 3.79 ± 0.43 |  |
| Fat-free mass (g) | 20.41 ± 0.28 | 20.50 ± 0.39 |  |
| Final: 14-Week HFHS |  |  |  |
| Body weight (g) | 38.95 ± 2.01 | 45.03 ± 1.31* |  |
| Fat mass (g) | 17.13 ± 1.56 | 22.45± 0.92* |  |
| Fat-free mass (g) | 21.82 ± 0.61 | 22.58 ± 0.46 |  |
| Energy Metabolism |  |  |  |
| Weekly food intake (g) | 16.25 ± 0.45 | 17.45 ± 0.32* |  |
| Weekly energy intake (kcal) | 76.86 ± 2.14 | 82.54 ± 1.51* |  |
| Weekly Total EE (kcal) | 62.38 ± 1.79 | 67.79 ± 1.39* |  |
| Weekly Resting EE (kcal) | 45.36 ± 1.59 | 51.06 ± 1.47* |  |
| Weekly Activity EE (kcal) | 10.30 ± 0.69 | 9.54 ± 0.59 |  |
| Respiratory Quotient | 0.83 ± 0.00 | 0.83 ± 0.01 |  |
| Weekly Cage Activity (m) | 696.78 ± 68.18 | 635.07 ± 31.51 |  |
| ANCOVA |  |  |  |
|  | SO | VCD |  |
| TEE (kcal, co-variate(s) BW, FFM+FM) | 64.23 ± 1.31 | 65.68 ± 1.41 |  |
| Co-variate Effect Size | Co-variate | p-value | Partial Eta<sup>2</sup> |
|  | BW | p<0.042 | 0.301 |
|  | FFM | p=0.160 | 0.171 |
|  | FM | p<0.006 | 0.502 |
| REE (kcal, co-variate(s) BW, FFM+FM) | 46.78 ± 1.41 | 49.43 ± 1.52 |  |
| Co-variate Effect Size | Co-variate | p-value | Partial Eta<sup>2</sup> |
|  | BW | p=0.216 | 0.125 |
|  | FFM | p=0.129 | 0.197 |
|  | FM | p<0.027 | 0.373 |
| Energy Intake (kcal, co-variate(s) BW, FFM+FM) | 79.41 ± 1.03 | 79.26 ± 1.11 |  |
| Co-variate Effect Size | Co-variate | p-value | Partial Eta<sup>2</sup> |
|  | BW | p<0.0001 | 0.723 |
|  | FFM | p=0.769 | 0.008 |
|  | FM | p<0.0003 | 0.706 |

Data are presented as mean ± standard error of the mean (SEM), n=7-8/group).

### VCD treatment induced severe steatosis following chronic HFHS diet

Next, we determined the effect of chronic HFHS feeding on hepatic steatosis and triglyceride levels in VCD mice. VCD treated mice showed substantially greater steatosis compared to the control group, consistent with the previous reports in OVX mice (Figure 7A)^16^. Steatosis severity, evaluated by percent score, was significantly higher in VCD livers (Figure 7B), while no difference in visual inflammation was observed in VCD livers compared to controls (Figure 7C). Also, no hepatocyte ballooning or fibrosis was observed in VCD livers compared to the controls (data not shown). These findings were associated with greater triglyceride levels in VCD livers compared to the control group (Figure 7D). However, we did not observe significant differences in gene expression of macrophage, inflammatory, and antioxidant markers between the two groups (Figure 7E).

**Figure 7.**
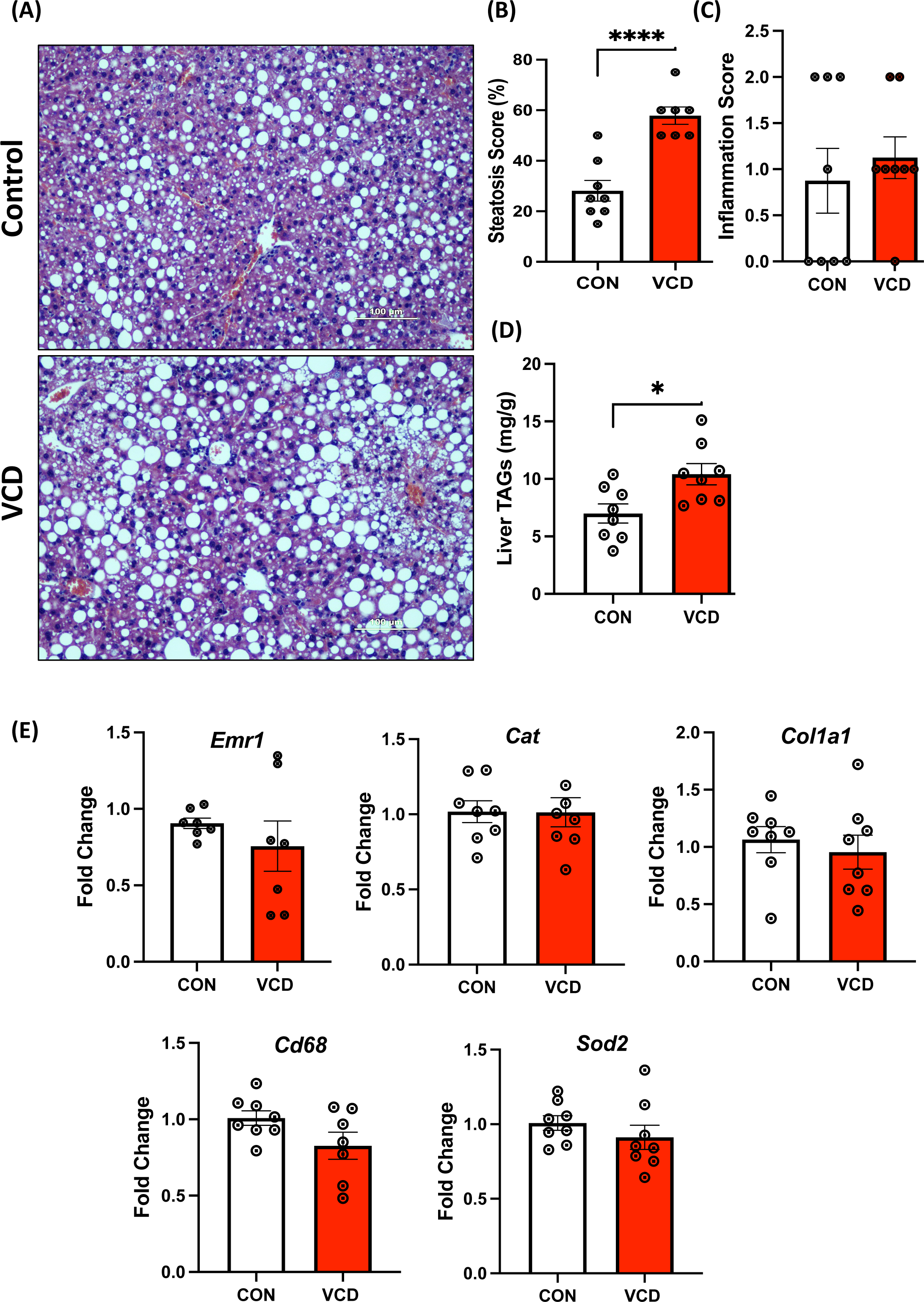
Experiment 3: Assessment of liver health and measures of key genes involved in liver health (Inflammation, oxidative stress) during 16-week HFHS in mice. (A) Representative H & E staining of liver sections showing more lipid droplets in the liver of VCD mice relative to control mice on 16-week of HFHS (scale bar: 100 μm), (B) Histological steatosis scoring (in percentage), (C) Inflammation score (0-none, 1-few, and 2-many), (D) VCD treated mice show increased liver triglyceride content, and (E) No major difference in expression of *Emr1, Cat, and Col1a1, Cd68, and Sod2* in VCD treated liver samples compared to control mice on 16 weeks HFHS diet. Data are presented as mean ± SEM (n=7-8/group), *p ≤ 0.05.

### VCD mice showed elevated ALT levels and distinctive cholesterol Trends on chronic HFHS diet

Similar to LFD and acute HFHS diet, we assessed the impact of VCD treatment on liver injury markers during chronic HFHS diet. Among the measured parameters, only the VCD treatment group following 16 weeks of HFHS, showed a significant increase in ALT levels, indicative of hepatocellular injury. Also, a notable upward trend in cholesterol levels was observed in the VCD-treated group compared to the control. No major changes were noted in ALP, AST, triglycerides, glucose, or HDL/LDL levels in response to VCD treatment during chronic HFHS diet. (Table 4).

**Table 4.**
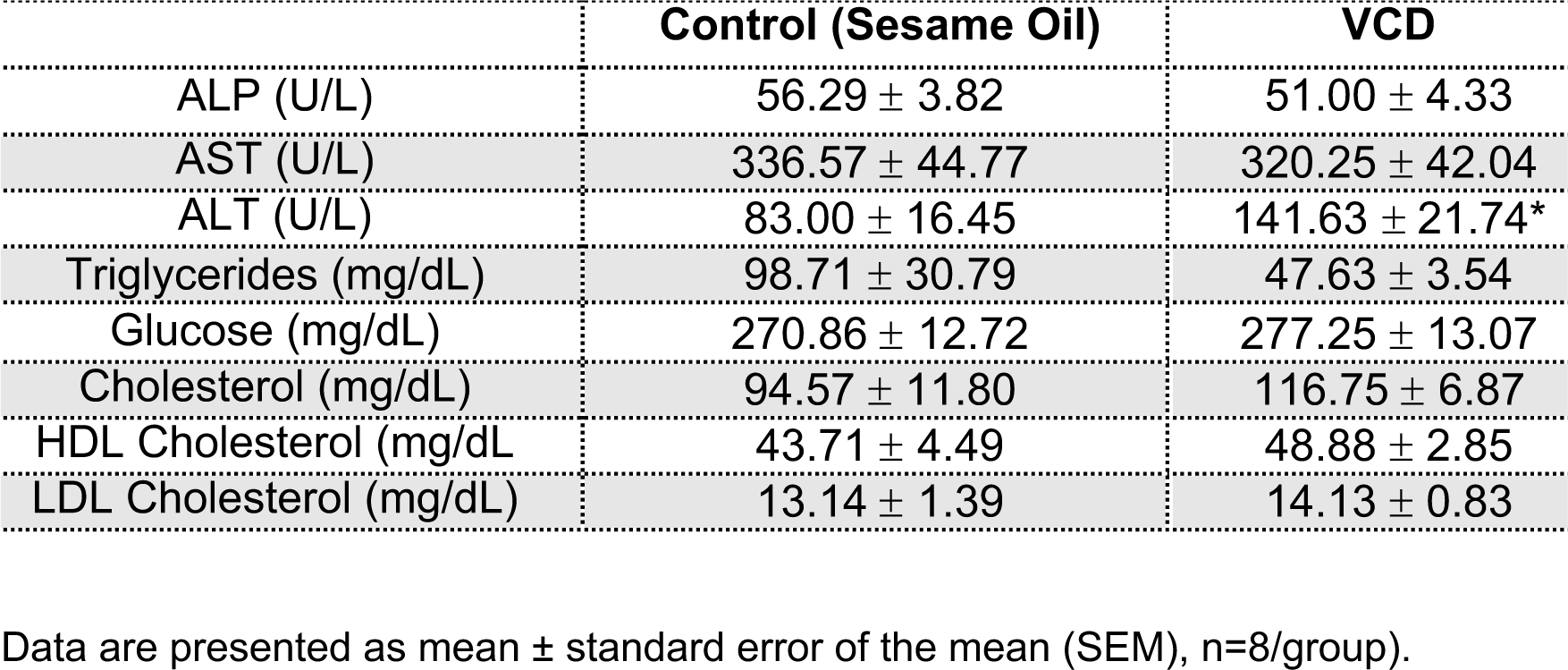
Experiment 3: Chronic high-fat/ high-sucrose diet (HFHS) serum cholesterol, glucose and liver injury markers

|  | **Control (Sesame Oil)** | **VCD** |
| --- | --- | --- |
| ALP (U/L) | 56.29 ± 3.82 | 51.00 ± 4.33 |
| AST (U/L) | 336.57 ± 44.77 | 320.25 ± 42.04 |
| ALT (U/L) | 83.00 ± 16.45 | 141.63 ± 21.74* |
| Triglycerides (mg/dL) | 98.71 ± 30.79 | 47.63 ± 3.54 |
| Glucose (mg/dL) | 270.86 ± 12.72 | 277.25 ± 13.07 |
| Cholesterol (mg/dL) | 94.57 ± 11.80 | 116.75 ± 6.87 |
| HDL Cholesterol (mg/dL) | 43.71 ± 4.49 | 48.88 ± 2.85 |
| LDL Cholesterol (mg/dL) | 13.14 ± 1.39 | 14.13 ± 0.83 |

Data are presented as mean ± standard error of the mean (SEM), n=8/group).

### Measures of hepatic mitochondrial respiration and oxidative stress

We next sought to determine whether liver mitochondrial respiratory phenotypes were different in VCD and control mice following a chronic HFHS diet. Isolated liver mitochondria respiratory capacity was measured using two different substrate combinations pyruvate/glutamate (PG) or lipids L-palmitoyl-CoA (PCoA)/L-palmitoyl-carnitine (PC). During PG respiration, we observed increased mitochondrial respiration at the basal and during State 3S in the VCD mitochondria compared to controls (Supplemental Figure 5A-C) with no major differences in mitochondrial coupling (CCR) and a higher trend in state3-basal respiration. Respiration of lipids (PC condition) was greater at all respiration states tested in isolated hepatic mitochondria from VCD-treated mice, except uncoupled (Figure 8A). Furthermore, CCR and ADP-dependent oxygen flux were significantly increased in VCD-treated hepatic mitochondria compared to the control group under these conditions (Figure 8B & C). To assess whether the increases in maximal lipid respiration represented differences in the ability of isolated mitochondria from chronic HFHS fed VCD mice to respond to changes in energy state, we performed a creatine kinase clamp (Figure 8D) ^45, 46^. The respiration of lipids was greater in VCD mice at all physiologically relevant ATP free energy states and the sensitivity of isolated mitochondria to increase respiration with increasing ATP free energy state (conductance, Figure 8E) was greater in the VCD mice compared to the control.

**Figure 8:**
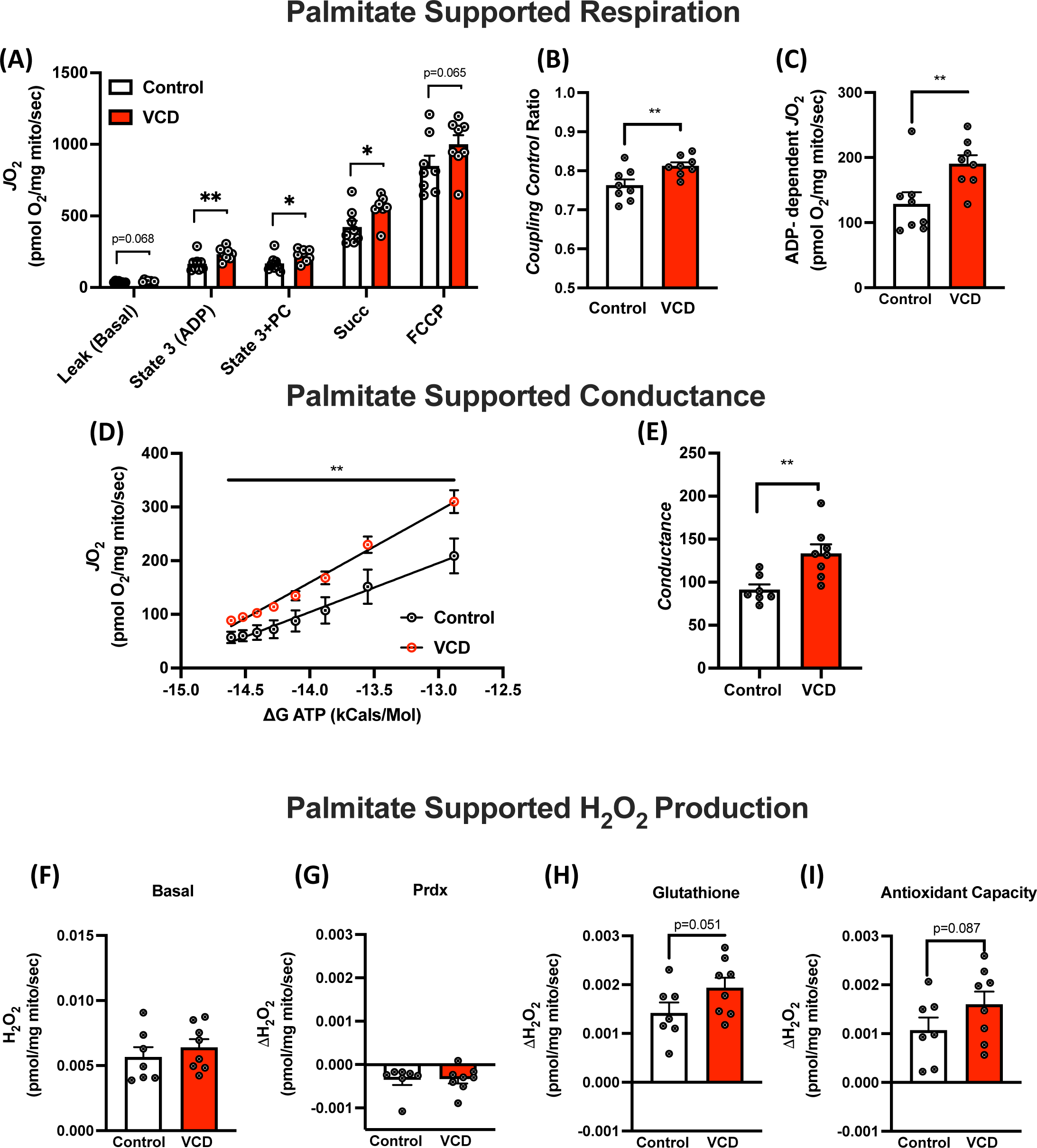
Experiment 3: Measures of hepatic mitochondrial respiratory capacity and H_2_O_2_ production of palmitate. (A) Using palmitate (palmitoyl-CoA & palmitoyl-carnitine) as a main substrate at basal, state 3 (ADP-dependent), Glutamine-dependent, succinate-dependent, cytochrome C-dependent, and FCCP uncoupled respiration. (B) The coupling control ratio was calculated by diving ADP-dependent respiration to the basal respiration. (C) ADP-dependent (State 3 – basal). Using the creatine kinase clamp, VCD mice have increased palmitate supported respiration at physiologically relevant ATP free energies (D) and (E) sensitivity of isolated mitochondria to changes in ATP free energy (conductance). (F) H2O2 emission during basal respiration of palmitate, (G) change due to peroxiredoxin antioxidant system inhibition, (H) change due to glutathione antioxidant system inhibition, and (I) maximal antioxidant capacity is presented. Respiration data from O2K were normalized using mitochondrial protein content by the BCA method. Data are presented as mean ± SEM (n=8/group), *p ≤ 0.05.

Interestingly, no differences were observed in basal H_2_O_2_ production during respiration of pyruvate or lipids (Supplemental Figure 5G-I & Figure 8F, respectively). However, while differences were observed in the antioxidant contribution of the peroxiredoxin system (Figure 8G), the glutathione antioxidant system activity was increased in isolated liver mitochondria from VCD mice on chronic HFHS compared to the control (Figure 8H). This difference helped generate a non-significant trend toward greater maximal H_2_O_2_ production in VCD mice liver mitochondria (Figure 8I). Together, these data demonstrate that VCD mice have increased adaptability of liver mitochondria to chronic HFHS feeding, however, this is accompanied by increased H_2_O_2_ generation and associated increases in antioxidant capacity.

## DISCUSSION

Through a series of experiments, we show that VCD-induced follicular atresia in the ovary leads to an expected loss of estrogen and increased FSH^7^, which is paired with changes in hepatic mitochondrial respiratory capacity and the hepatic mitochondrial proteome. Further, bulk RNA sequencing in the liver revealed that VCD livers display increased lipid/cholesterol biosynthesis pathway gene transcription. However, these alterations were not linked to significant differences in hepatic lipid storage, adiposity, or body mass during LFD feeding. Follow-up short-term and chronic HFHS studies investigated how VCD treatment influenced whole-body metabolic adaptations and susceptibility to weight gain and MASLD onset. We found that VCD mice have a greater preference for lipid utilization and possibly impaired metabolic flexibility on LFD, with no impact on energy intake and diet-induced changes in body weight or body composition. However, both acute and chronic HFHS feeding resulted in increased hepatic lipid accumulation in VCD mice compared to controls. Further, VCD mice chronically fed HFHS gained significantly more weight and adiposity than controls. Overall, these results point to a subtle whole-body metabolic phenotype induced by VCD that is more potently elicited by chronic HFHS feeding.

The prevalence of metabolic diseases, including MASLD, significantly increases after menopause^9^. Previous findings have shown that pre-menopausal women and female rodents exhibit remarkable resistance to diet-induced metabolic and mitochondrial dysfunction compared to age-matched males ^47, 48^. This sexual dimorphism is believed to be mediated, at least partially, by the ovarian hormone estrogen. However, as estrogen levels fall during menopause, females experience increased susceptibility to weight gain and metabolic disease onset^49^. Several studies have demonstrated that reproductive hormones (e.g., estrogen and FSH) act as key regulators of systemic and liver-specific energy metabolism, and alterations in these hormones with menopause negatively impact metabolic homeostasis^21^. Specifically, loss of estrogen signaling through its cognate receptors (e.g, ERα, ERβ, and GPER) plays a crucial role in the menopause phenotype^15^. ERα primarily mediates estrogen’s protective role in maintaining energy expenditure, food intake, glucose production, and insulin resistance in metabolic tissues (liver, adipose and muscle) and the hypothalamus. We recently reported that liver-specific ERα knockout mice with normal estrous cycling do not show a liver mitochondrial phenotype or increased susceptibility for steatosis compared to controls ^50^. However, estrogen depletion in ovariectomized mice (OVX) results in a pronounced metabolic phenotype, including increased appetite, lower physical activity, increased adiposity, impaired mitochondrial function in the liver, and steatosis, which can be reversed with estrogen replacement therapy ^16, 51^. Previous results do suggest that ERα is necessary for estrogen treatment of hepatic steatosis in mice that have lost ovarian function ^52^. Additionally, estrogen protects against high-fat diet-induced insulin resistance and glucose intolerance in the liver, and its decline during menopause may lead to hepatic fat accumulation ^16, 51, 53–58^. In addition, blocking FSH action has been shown to impact body weight and fat mass in female mice by reducing adiposity and increasing thermogenesis and adipose mitochondrial density, thus influencing energy expenditure^59–61^.

Loss of estrogen following menopause or ovariectomy is associated with increased body weight and adiposity ^51, 54–58^, which is associated with decreased total EE and increased energy intake. In this study we did not observe significant differences in body weight, body composition, total EE, or energy intake between the control and VCD-treated groups during a LFD nor in response to a 1-week HFHS diet. This suggests that the onset of a menopause-like phenotype induced by VCD treatment does not immediately affect systemic energy homeostasis in mice. However, chronic HFHS feeding resulted in greater body weight and adiposity increases in VCD compared to control mice. These findings are similar to previously reported literature in OVX mice on HFHS diets ^62, 63^.

This collection of experiments allowed for observing changes in systemic energy metabolism across LFD, acute, and chronic HFHS. While energy intake and expenditure differences were limited to diet-induced reductions in cage activity and greater reduction in activity EE, VCD mice demonstrated lower RQ during both LFD and acute HFHS. The preference for fat utilization in VCD mice at baseline and a potentially reduced capacity to adapt to changes in substrate availability during the HFHS diets suggest disrupted metabolic homeostasis and a potential propensity for weight gain ^64^. Whole body metabolic flexibility is linked to fuel metabolism in various tissues (liver, muscle, and adipose) and has been proposed to be causative in the development of impaired insulin action (lower glucose utilization in muscle and elevated hepatic glucose output), dysregulation of lipolysis in adipose, and altered fat oxidation in liver and muscle. ^65^. It is also linked to ectopic lipid storage in metabolic organs including the liver. Thus, preference for lipid utilization in VCD mice may be an early signal of metabolic dysregulation that deserves further study. Importantly, chronic HFHS feeding resulted in increased measures of systemic energy metabolism in the VCD mice compared to the control. However, further data analysis of the data to assess the impact of the differences in body weight and composition demonstrated that all differences in energy intake and expenditure were due to the larger mass of the VCD mice. Interestingly, fat mass, and not fat-free mass, was found to be a powerful co-variate impacting systemic energy metabolism. We have previously observed this in another unrelated experiment involving chronic HFHS feeding of normally cycling female mice ^66^. Combined, these findings are interesting considering the low metabolic rate of adipose tissue and the numerous observations of a high correlation of FFM with energy expenditure, particularly resting EE ^67^. However, differences in body composition have been observed to alter the relationship to energy expenditure ^68, 69^, and are further complicated by the difference in fat mass and fat-free mass composition and distribution in males and females ^70^. Further investigation is needed into how the relationship of body composition to energy expenditure changes during chronic energy dense diet feeding in females compared to males.

Previously, we have demonstrated the role of impaired mitochondrial respiratory phenotypes in the sexually dimorphic development of hepatic steatosis ^27, 38, 65^. This work demonstrated that female mice have a much different hepatic mitochondrial phenotype compared to males including lower basal respiration, greater coupling control ratio, and lower H_2_O_2_ emission ^27, 65^. Moreover, female mitochondria display a robust capacity to up-regulate respiration and maintain a coupling control ratio on a HFHS or exercise plus a HFHS diet, a response that does not occur in male hepatic mitochondria ^38^. However, we and others have also shown that OVX dramatically diminishes the female hepatic mitochondrial phenotype. ^16^. Thus, we assessed the mitochondrial phenotype of female mice treated with VCD to determine whether loss of follicular function impacts hepatic mitochondrial respiratory capacity. Hepatic mitochondria from VCD mice on a LFD displayed reduced respiration of lipid substrate at all respiratory states. Unfortunately, no measurement of H_2_O_2_ emission was performed nor did we test different substrates (pyruvate vs. lipid) in these studies. However, these results suggest that the VCD-induced induction of a menopause-like phenotype significantly changes the hepatic mitochondrial respiratory capacity in the direction of compromised fat oxidation in liver mitochondria. Also, these reductions in mitochondrial respiration could set the stage for the development of steatosis over time or if pushed with an energy dense, HFHS diet.

Additionally, the impact of chronic HFHS feeding on mitochondrial respiratory phenotypes was assessed in both VCD and control livers. In these experiments, we performed respiration with both pyruvate and lipid-based substrates and quantified H_2_O_2_ emission. Previous results in humans and mice have suggested that a HFHS diet or the presence of steatosis or obesity leads to compensatory upregulation of hepatic mitochondrial respiratory capacity ^21, 22, 71^. Interestingly, we found that VCD mice displayed a similar upregulated compensation of respiratory function when compared to female controls and this occurred with both pyruvate and lipid substrates and across different respiration states (basal, State3, and State3S). Also, the increased VCD mitochondrial respiration of lipids was observed at physiologically relevant ATP-free energy levels and demonstrated greater sensitivity to the change in energy state. Further, these increases in oxygen consumption were accompanied by increases in total antioxidant capacity. Thus, despite baseline deficits in mitochondrial respiratory capacity shown on a LFD, the VCD mice display evidence of a more significant compensatory increase in respiratory capacity on a HFHS diet that induces steatosis than controls. Overall, these results suggested that the VCD displayed a greater compensation than control but still displayed greater steatosis. The field is gradually evolving its understanding of hepatic mitochondrial oxidative capacity and steatosis risk to understand that mitochondrial compensation is an initial adjustment, but that as obesity and associated changes in steatosis and liver injury worsen, mitochondrial adjustments switch from compensatory increases to a decline in function. This has been reported previously in two human studies in which liver mitochondrial phenotypes were captured with respiratory and fat oxidation outcomes ^72, 73^. A possible interpretation is that more significant upregulation of mitochondrial respiratory function in VCD mice paired with greater hepatic lipid storage may be a short-term adjustment that is linked to a greater overload of hepatic metabolism than in controls. Moreover, it is possible that this upregulation in respiration in VCD may lead to a faster decline in mitochondrial function as liver injury develops. More work on female liver mitochondrial physiology is needed to uncover its potential role in etiology of metabolic disease.

The deficits in hepatic mitochondrial respiratory capacity observed in LFD fed VCD mice correspond with the observed global reductions in proteins associated with the electron transport system (ETS), likely contributing to the decreased capacity for oxidative phosphorylation. While the effects of estrogen on hepatic mitochondrial function remain largely unknown, estrogen has been shown to directly exert transcriptional regulation over several proteins involved in oxidative phosphorylation, particularly through estrogen receptor α (ERα) signaling ^74^. Additionally, evidence in skeletal muscle has suggested that estrogen mediates the activation of DNM1L/Drp1, a critical protein necessary for mitochondrial fission and activation of mitophagy ^75^. DNML1/Drp1 was recently observed to be critical in improving diet-induced steatohepatitis, via reduced ER stress and improved mitochondrial respiratory capacity ^76^. The observed decrease in DNM1L/Drp1 and BNIP3 in VCD mice supports the potential role of estrogen in maintaining hepatocyte mitophagy pathways. Further, these reductions in fission/mitophagy proteins may account for the increased adaptability in mitochondrial respiratory capacity present in chronic HFD feeding in VCD mice. A recent study has shown that in an obese state, the knockdown of DNM1L/Drp1 can promote improvements in systemic metabolism by promoting the formation of enlarged mitochondria that possess the expanded respiratory capacity and reduced H_2_O_2_ production ^77^ however, this upregulation may only be beneficial for a certain amount of time and lead to longer term pathologies which have recently been reported in liver-specific DNML1/Drp1 ko mice which showed more pathological liver injury after chronic feeding ^78^. The reduction of DNM1L/Drp1 in VCD mice may promote an imbalance in mitochondrial fission/fusion events, resulting in enlargement of the hepatic mitochondrial morphology which may support greater functional capacity within the liver which was seen after HFHS ^79^. Additionally, as stated, evidence in rodents and humans supports acute increases in hepatic mitochondrial oxidative capacity and tricarboxylic acid (TCA) cycle flux in individuals with MASLD prior to the development of mitochondrial dysfunction^19, 21, 80^. This compensatory increase in oxidative metabolism and TCA cycle flux in combination with enlarged mitochondrial morphology may explain the increased mitochondrial respiratory capacity seen in VCD mice following HFHS feeding.

We also performed bulk RNA seq analysis of livers from VCD and control mice on a LFD to determine how the induction of a menopause-like phenotype may influence liver transcriptional regulation and metabolic pathways. Our bulk liver RNA seq results revealed a significant number of differentially expressed genes (190) in the livers of VCD mice, with a dramatic upregulation in the expression of genes associated with *de novo* lipogenesis and cholesterol synthesis pathways before the onset of steatosis development. Our data suggest distinctive transcriptional dysregulation in VCD livers, particularly in the context of lipogenic and cholesterol synthesis pathways, establishing an initial deficit in liver lipid metabolism that may increase susceptibility to steatosis, especially under high fat/sucrose diets. In postmenopausal women, the decline in estrogen levels has been shown to interact with weight gain and fat mass and progressive transition to higher rates of steatosis (ref). However, it remains unknown whether the decline of estrogen, an increase in FSH, or changes in other hormone levels induce the steatosis phenotype ^24, 81, 82^. Regardless of the current gaps in our knowledge of how menopause is linked to steatosis, the VCD model demonstrated increased steatosis during short-term HFHS and chronic HFHS feeding compared to controls. Histological analysis confirmed the presence of steatosis in both groups, with VCD-treated mice exhibiting more abundant lipid droplets. Furthermore, VCD mice exhibited increased liver triglyceride levels compared to control, with a greater difference observed after prolonged exposure to the HFHS diet. Overall, these results indicate that VCD treated mice represent a valuable model system to study how the loss of follicular estrogen production with an increase in FSH increases susceptibility to diet-induced steatosis.

One potential limitation of our study is the absence of mitochondrial outcomes in the acute HFHS condition, which would have allowed us to determine how VCD hepatic mitochondria adjust to short-term energy surplus provided by the HFHS diet. Further, a lack of a control LFD group within our chronic HFHS diet limits the assessment of the prolonged menopause phenotype independent of diet. Additionally, these experiments utilized the C57Bl6/J mouse strain. While this strain is widely used in metabolic studies, it has inherent genetic defects predisposing it to greater diet-induced metabolic disease development. Current studies are ongoing to compare chronic LFD vs. HFHS in controls and VCD mice. Additionally, we performed VCD in relatively young mice and not in older mice that would track with a human relative age in which menopause occurs.

In conclusion, our examination of the VCD model revealed a subtle whole-body metabolic phenotype that required a HFHS diet to reveal increases in adiposity and significantly increased hepatic steatosis compared to control mice. However, livers from LFD-fed VCD mice did display significant differences in global gene expression in the liver, particularly for cholesterol synthesis genes. Moreover, there were significant differences in hepatic mitochondrial proteome and respiratory function compared to controls. These outcomes demonstrate the powerful role that loss of ovarian follicular function, particularly a loss of estrogen, and upregulation of FSH has on the programming of hepatic transcriptional regulation and the mitochondrial proteome. Future studies are needed to define the VCD model as a rigorous model for the study of the increased risk for metabolic disease development in women after menopause.

## Acknowledgments

The authors would like to thank Dr. Lane Christenson’s and Hong Xiaoman for providing training on mouse staging techniques and dissecting ovary in mice and KUMC Core seq facility for generating RNA sequencing data used in this study.

## Author Contributions

EMM, JPT and RK Conceptualized and designed the experiments; RK, MP and JP performed experiments; RK, EF, MO, MS, KS and EMM analyzed data and interpreted results of experiments; RK prepared original manuscript; EMM, AJL, EF, KS and JPT edited and revised manuscript. Supervision & Funding Acquisition: JPT and EMM.

---

**Supplementary** Figure 1: (A) Cytological assessment, (B) Representative H & E staining of liver sections in the VCD mice and control mice on LFD in experiment 1, and (C) serum FSH levels for experiment 2. Data are presented as mean ± SEM (n=8/group), *p ≤ 0.05.

**Supplementary** Figure 2**: Experiment 1: VCD treatment on LFD induced the expression of lipogenic and cholesterol synthesis and downregulated the pathway involved in cytokines regulation in liver homogenate.** (A) Pathways downregulated in VCD (n=4/group). (B) RNA Seq in VCD livers showing higher expression of *Fasn, Elvol6, Scd1, Acaca, Acacb, Acly and Cyp8b1* in VCD livers compared to control mice, (n=4/group), * indicates Unadjusted p-values from limma analysis (*p ≤ 0.05).

**Supplementary** Figure 3**: Experiment 1:** Boxplots showing liver mRNA level of cholesterol transport genes in control and VCD-treated mice on LFD (n=4/group, *p ≤ 0.05).

**Supplementary** Figure 4**: Experiment 1:** Proteomics analysis in VCD hepatic mitochondria showing (A) Mitocarta proteins upregulated or downregulated with statistically significant p-values. (B) Mitocarta proteins upregulated or downregulated with LogFC: ± 1 with statistically significant p-values, (C) Reactome Pathway analysis shows metabolism is one of the most enriching classes with protein set cross-referenced to Mitocarta, and (D) Protein network for metabolic pathway enriched in VCD. Data are presented as mean ± SEM (n=8/group), *p ≤ 0.05.

**Supplementary** Figure 5**: Experiment 3: Measures of hepatic mitochondrial respiratory capacity and H_2_O_2_ production.** Mitochondrial respiratory capacity was measured using pyruvate/glutamate (PG) as a main substrate at basal, state 3 (ADP-dependent), Glutamine-dependent, succinate-dependent, cytochrome C-dependent, and FCCP uncoupled respiration. (B) The coupling control ratio was calculated by diving ADP-dependent respiration to the basal respiration. (C) State 3 (ADP-dependent) - basal oxygen flux. (D-F) Pyruvate- dependent H_2_O_2_ emission. Respiration data from O2K were normalized using mitochondrial protein content by BCA method. Data are presented as mean ± SEM (n=8/group), *p ≤ 0.05.

