## Supplemental Figures for "VCD-induced menopause mouse model reveals reprogramming of hepatic metabolism"

### Supplementary Figure 1

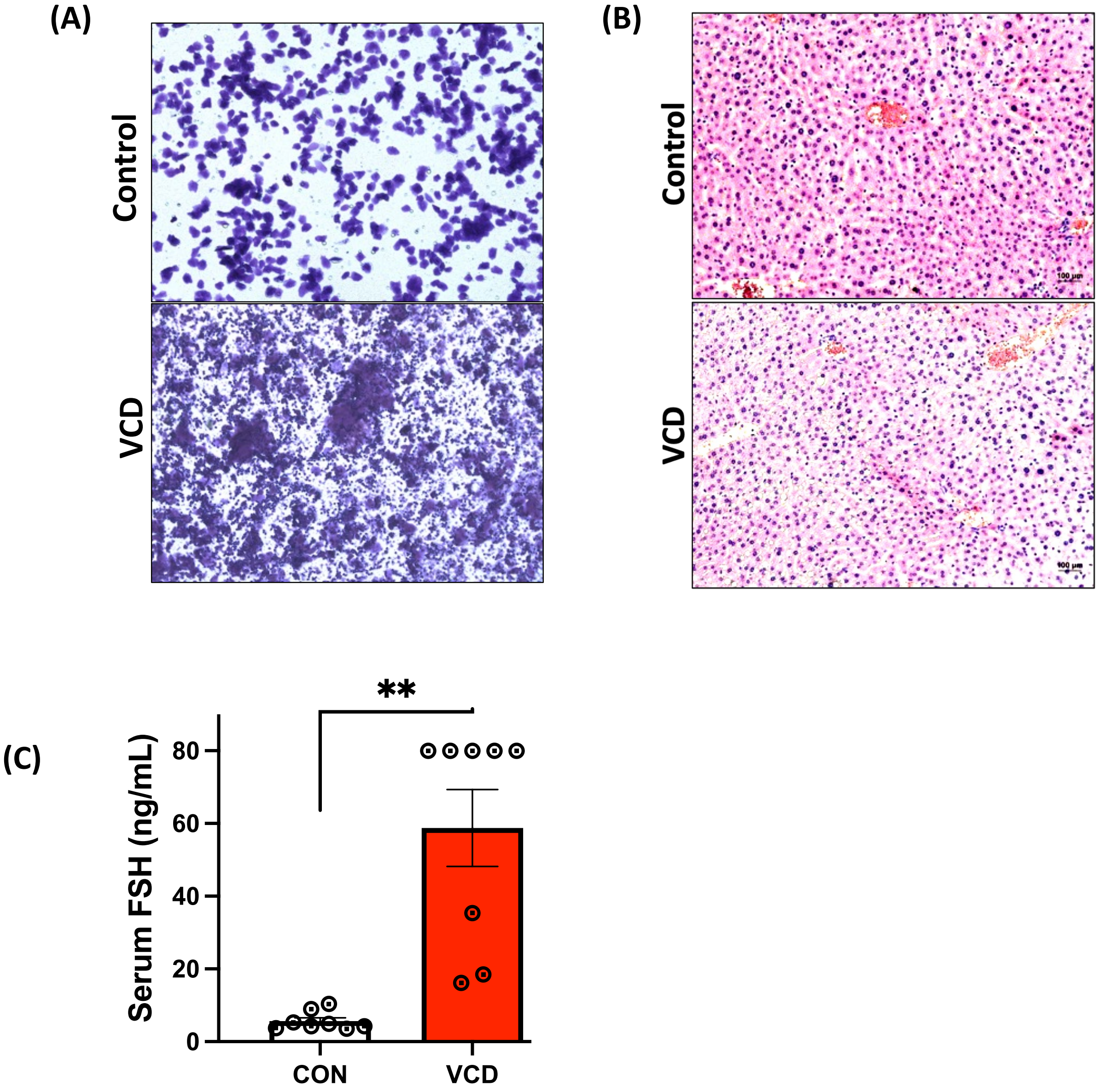

### Supplementary Figure 2

(A)

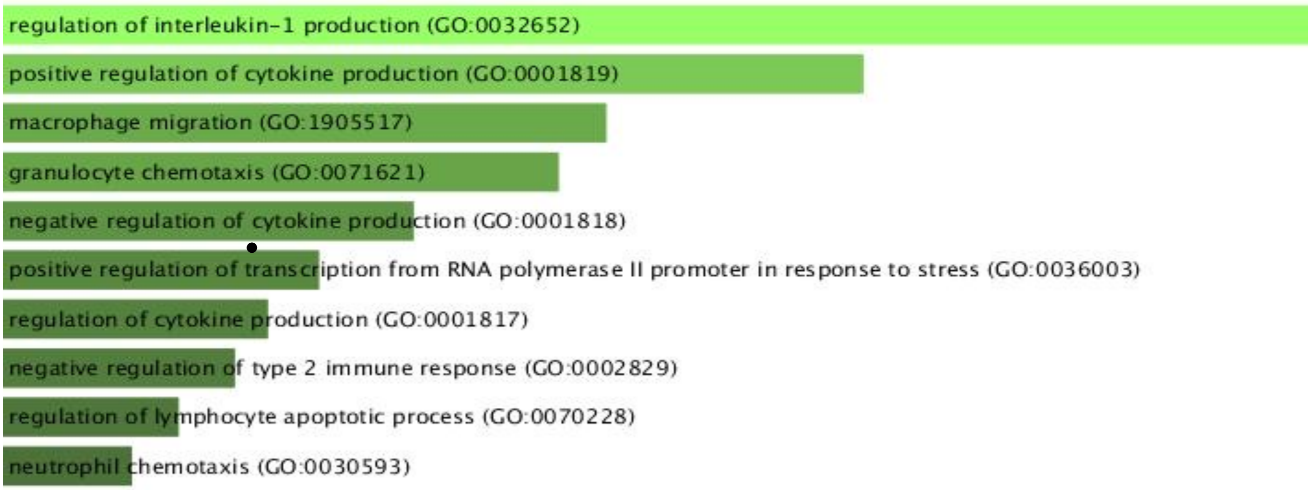


(B)

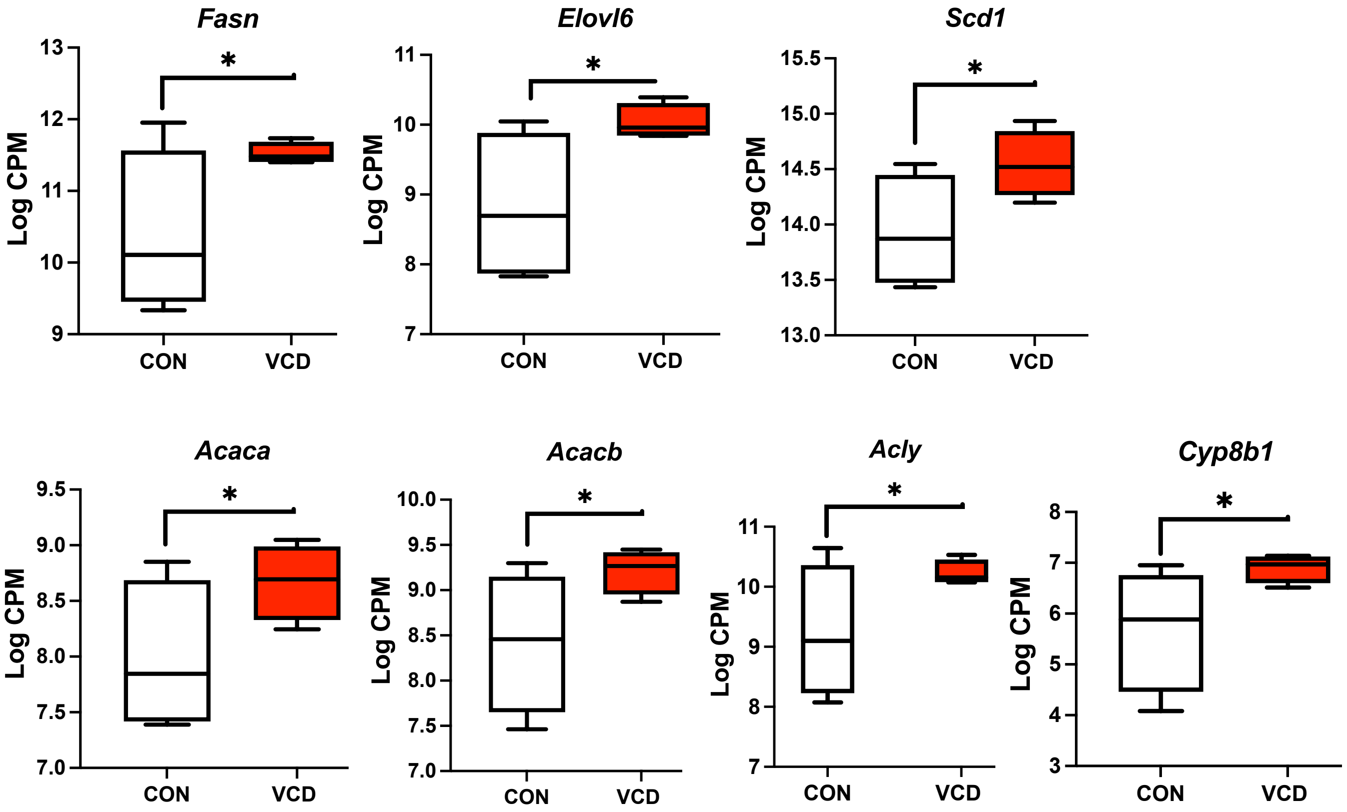

### Supplementary Figure 3

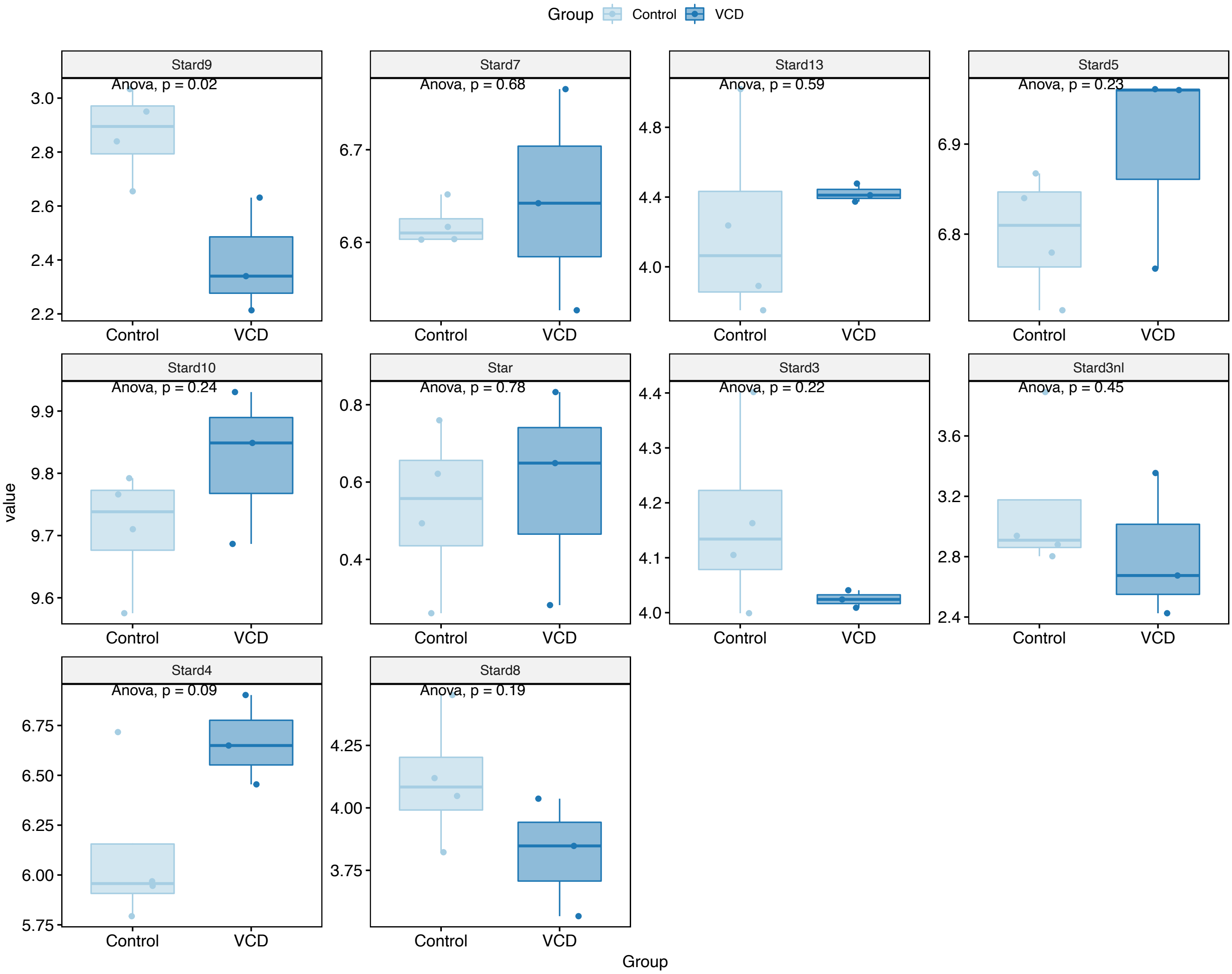

### Supplementary Figure 4

(A)

**Proteins cross referenced with mitocarta**

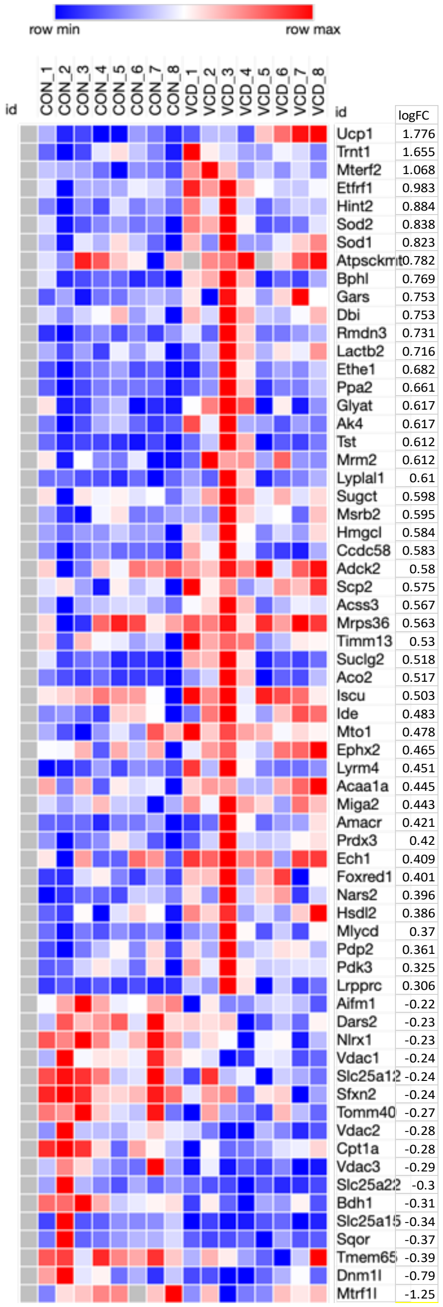


(B)

**Proteins cross referenced with mitocarta (with LogFC cutoff ± 1)**

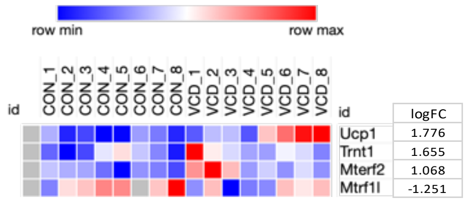


(C)

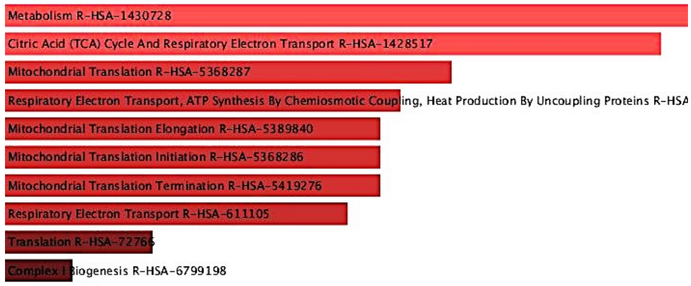


(D)

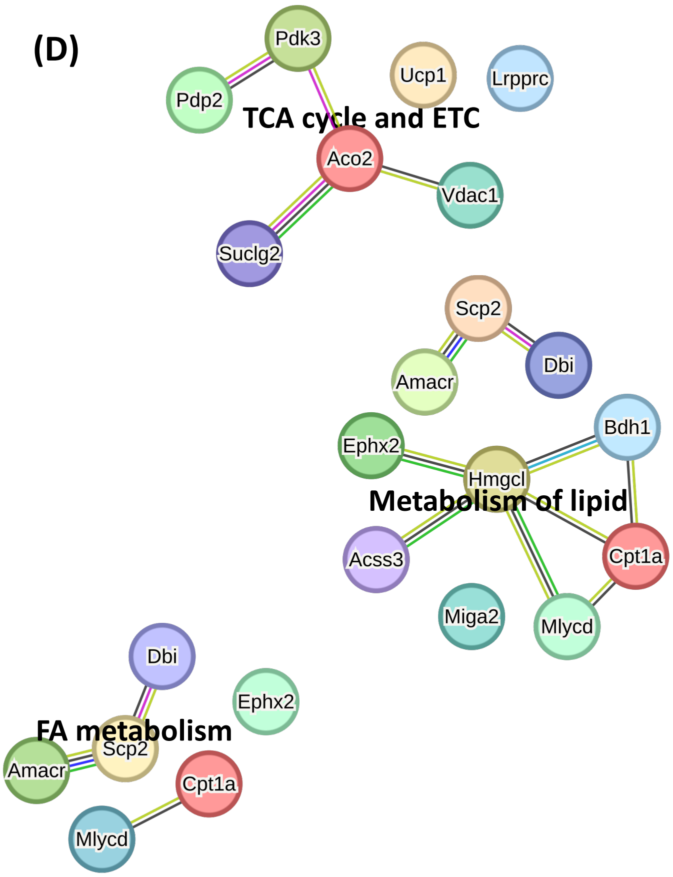

### Supplementary Figure 5

#### Pyruvate Respiration

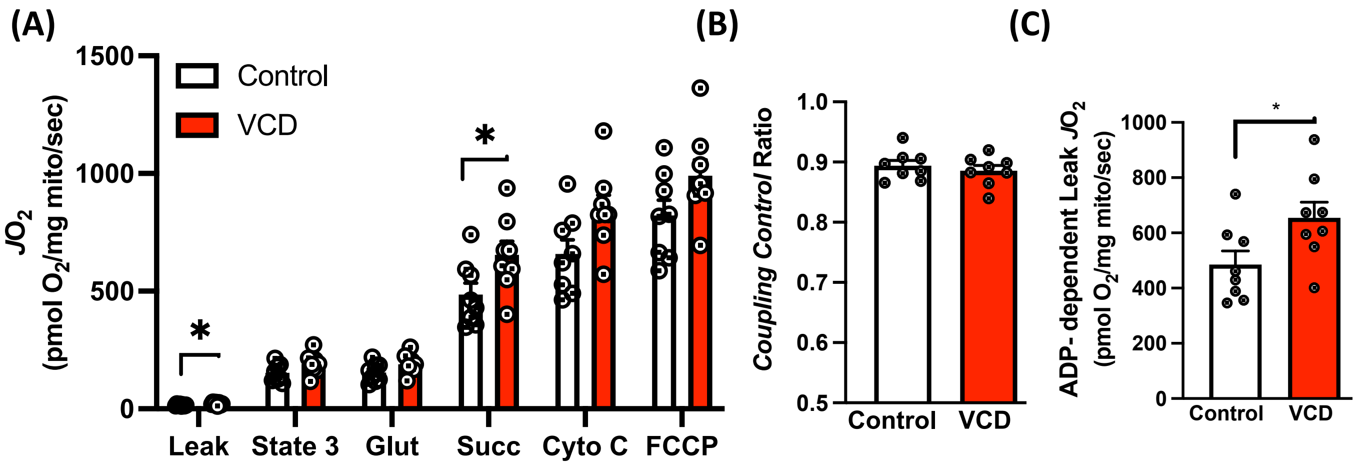


#### Pyruvate $H_2O_2$ Emission

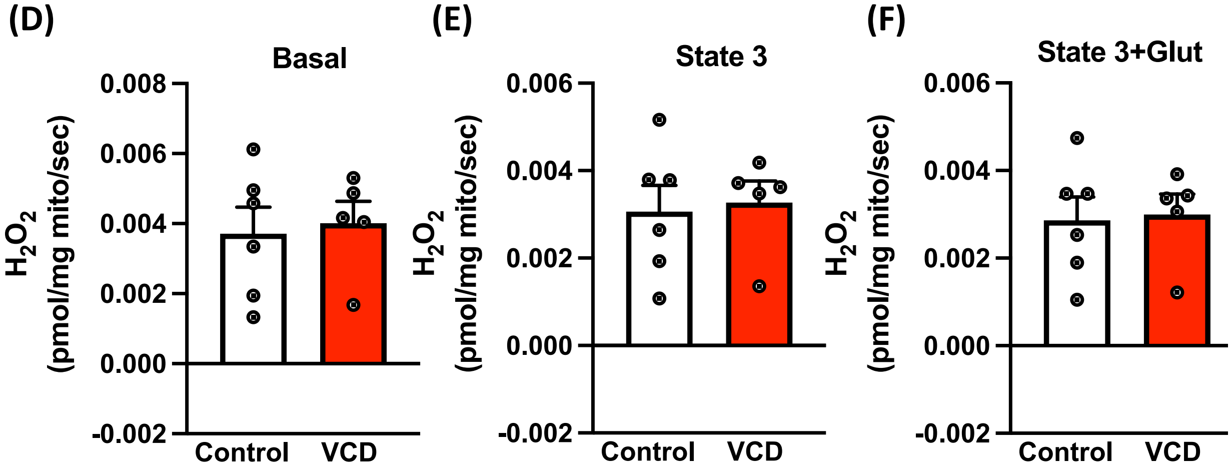

### Supplementary Table 1

| | Control | VCD |
|---|---|---|
| ALP (U/L) | 85.4 ± 24.71 | 71.00 ± 7.85 |
| AST (U/L) | 234.60 ± 39.76 | 238.17 ± 56.52 |
| ALT (U/L) | 44.80 ± 18.46 | 34.50 ± 8.96 |
| Triglycerides (mg/dL) | 37.60 ± 3.25 | 44.00 ± 4.61 |
| Glucose (mg/dL) | 188.40 ± 25.09 | 252.00 ± 31.45* |
| Cholesterol (mg/dL) | 53.60 ± 8.94 | 57.50 ± 6.72 |
| HDL Cholesterol (mg/dL | 27.60 ± 4.57 | 30.00 ± 4.46 |
| LDL Cholesterol (mg/dL) | 7.40 ± 0.51 | 8.00 ± 0.63 |

**Table 1. Experiment 1: LFD serum cholesterol, glucose, and liver injury markers**
Data are presented as mean ± standard error of the mean (SEM), n=8/group).

### Supplementary Table 2

| | Control | VCD |
|---|---|---|
| ALP (U/L) | 55.67 ± 4.13 | 65.17 ± 4.59 |
| AST (U/L) | 117.50 ± 22.33 | 174.17 ± 26.45 |
| ALT (U/L) | 54.50 ± 17.45 | 80.00 ± 14.20 |
| Triglycerides (mg/dL) | 38.50 ± 4.19 | 36.67 ± 1.38 |
| Glucose (mg/dL) | 236.50 ± 14.38 | 232.50 ± 21.61 |
| Cholesterol (mg/dL) | 104.67 ± 12.46 | 119.50 ± 8.31 |
| HDL Cholesterol (mg/dL | 45.83 ± 4.42 | 45.67 ± 1.84 |
| LDL Cholesterol (mg/dL) | 13.33 ± 0.67 | 14.33 ± 1.23 |

**Table 2. Experiment 2: LFD serum cholesterol, glucose, and liver injury markers**
Data are presented as mean ± standard error of the mean (SEM), n=8/group).
